# The spatiotemporal structure of neural activity in motor cortex during reaching

**DOI:** 10.1101/2025.10.17.683171

**Authors:** Ryan A. Canfield, Tomohiro Ouchi, Hao Fang, Beatrice Macagno, Lydia I. Smith, Leo R. Scholl, Amy L. Orsborn

## Abstract

Intracortical brain-computer interfaces (BCI) leverage knowledge about neural representations to translate movement-related neural activity into actions. BCI implants have targeted broad cortical regions known to have relevant motor representations, but emerging technologies will allow flexible targeting to specific neural populations. The structure of motor representations in neural populations across frontal motor cortices, which span centimeters, has not been well characterized. Here, we investigate how motor representations and population dynamics (temporal coordination) vary across a large expanse of frontal motor cortices. We used high-density, laminar, microelectrode arrays to record many neurons and then sampled neural populations across frontal motor cortex in two monkeys while they performed a reaching task. Our experiments allowed us to map neuronal activity across three spatial dimensions and relate them to movement. Target decoding analysis revealed that target direction information (one key aspect of task information) was heterogeneously distributed across the cortical surface and in depth. Similarly, we found that the temporal dynamics of different neural populations was highly variable, but that the amount of task information predicted which neural populations had similar dynamics. The neural populations with the most similar dynamics were composed of neurons with high task information regardless of spatial location. Our results highlight the spatiotemporal complexity of motor representations across frontal motor cortex at the level of neurons and neural populations, where well-learned movements consistently recruit a spatially distributed subset of neurons. Further insights into the spatiotemporal structure of neural activity patterns across frontal motor cortex will be critical to guide future implants for improved BCI performance.

**Significance Statement:** Motor brain-computer interfaces (BCI) translate neural activity into movement, but how to target implants within motor cortices to maximize performance remains unclear. We used high-density recordings of neural activity spanning a large cortical area and related them to movement to map the spatial distribution of task information and the evolution of neural population activity over time. Our measurements revealed that neurons with the most task information were highly distributed across cortex yet also evolved coherently in time, suggesting that spatially distributed neurons coordinate to control movements. Our results provide new links between neuron- and population-level maps of motor representations, and highlight the complex spatiotemporal structure of activity that may need to be considered when designing next-generation BCIs.

## Introduction

Intracortical brain-computer interfaces (BCI) translate neural activity into movement and provide promising therapies for motor rehabilitation (Hochberg et al., 2006; Collinger et al., 2013; Silversmith et al., 2020). Current BCIs leverage existing knowledge about the structure of movement information in the brain to target implants (Dadarlat et al., 2023), often targeting areas where electrical stimulation evokes movement or using functional MRI mapping. These mapping approaches can guide targeting on the scale of several millimeters (Park et al., 2001; Kunigk et al., 2024), which approximately matches the spatial resolution of common implant technologies like multi-electrode arrays with fixed, millimeter-scale geometries (e.g., Utah Arrays are often 4 x 4 mm). New technologies now allow flexible probe targeting to specific neurons and neuronal populations (Jun et al., 2017; Musk and Neuralink, 2019; Yang et al., 2019). The spatiotemporal structure of single neuron and population activity in frontal motor cortices, however, has not been well-characterized, limiting our ability to leverage these technologies for BCIs.

Task information representations in frontal motor cortices appear to be both spatially and temporally complex. Many studies found that the cortical areas where electrical stimulation evoked movements did not have clearly defined spatial boundaries (Asanuma and Rosén, 1972; Park et al., 2001; Chehade and Gharbawie, 2023). For instance, areas that evoked finger movements were intermixed with areas that evoked wrist movement, consistent with anatomical projection tracing that suggests small areas in motor cortex have clustered intracortical connections (Huntley and Jones, 1991). Studies that map relationships between neural activity and behavior at multi-neuron resolution further show that task information is unevenly distributed across frontal motor cortical locations and that neural activity patterns from neighboring locations evolve differently in time (Chehade and Gharbawie, 2023; Ouchi et al., 2025). Single-unit activity during movement also shows a wide range of temporal relationships to reaching (Churchland and Shenoy, 2007), and temporal relationships between individual neurons and behavior suggest they may coordinate via functionally defined networks (Peters et al., 2014; Moore et al., 2024). However, most studies of motor representations measured neuronal populations across limited regions of cortex (Churchland and Shenoy, 2007; Peters et al., 2014; Moore et al., 2024) or mapped individual neuron properties across large cortical areas (Schieber and Hibbard, 1993; Georgopoulos et al., 2007), leaving open the question of how task-relevant information is temporally structured across neural populations in frontal motor cortex.

Understanding the spatiotemporal structure of task representations across motor cortices will likely benefit from neural population analyses, which recently revealed distinct temporal structure in neural activity patterns during reaching (Churchland et al., 2012; Gallego et al., 2018, 2020; Lara et al., 2018). Existing population analyses, however, pooled neural activity across large areas within and sometimes across multiple motor cortical regions (Churchland et al., 2012; Gallego et al., 2020). Therefore, the spatial structure of population coordination during movement production remains unclear.

Here we examine how target-direction information, one aspect of task information, is spatiotemporally organized across the motor cortical surface and depth during reaching. We used a high-density, laminar, micro-electrode array to make spatially precise single-unit recordings from frontal motor cortical neural populations as monkeys performed a well-learned reaching task. These populations span the cortical surface from dorsal pre-motor cortex (PMd) to the putative arm region of primary motor cortex (M1), and cortical depths of nearly 4mm.

Population decoding revealed that task information was unevenly distributed in depth and across the cortical surface. Analyzing population dynamics also showed that the temporal structure of neural populations varied with minimal relationship to spatial location. While spatial position was only marginally predictive of which neural populations had similar temporal structure, the task information represented within the population was strongly predictive. We used task information to define neural populations and found that similar dynamics were primarily driven by a subset of units with high task information rather than spatial location. Linking neural population dynamics to task information further suggested that these coordinated dynamics contained similar task representations. Our results reveal new details of the spatiotemporal structure in frontal motor cortical representations consistent with emerging ideas of task-specific, spatially distributed functional networks. Our results suggest, in turn, that improving BCIs may require pairing advanced probe technologies with improved functional mapping.

## Materials and Methods

### Surgical procedures

All procedures were conducted in compliance with the NIH Guide for the Care and Use of Laboratory Animals and were approved by the Institutional Animal Care and Use Committee at the University of Washington. Two male rhesus macaques (*Macaca mulatta*) aged 10 years old (Monkey 1) and 11 years old (Monkey 2) were implanted with a multi-modal chamber stereotaxically targeted over left frontal motor cortex (Fig. 1C), using surgical procedures fully described in (Ouchi et al., 2025). The chamber leveraged a semi-chronic artificial dura (AD) that allowed visual and electrophysiological access to a 19 mm diameter window that included premotor cortex and primary motor cortex. A semi-chronic grid with Ø1 mm holes was rigidly attached to the chamber and implanted over the AD to stabilize cortical movements and allow spatially precise intracortical recordings to repeatable locations.

**Figure 1.**
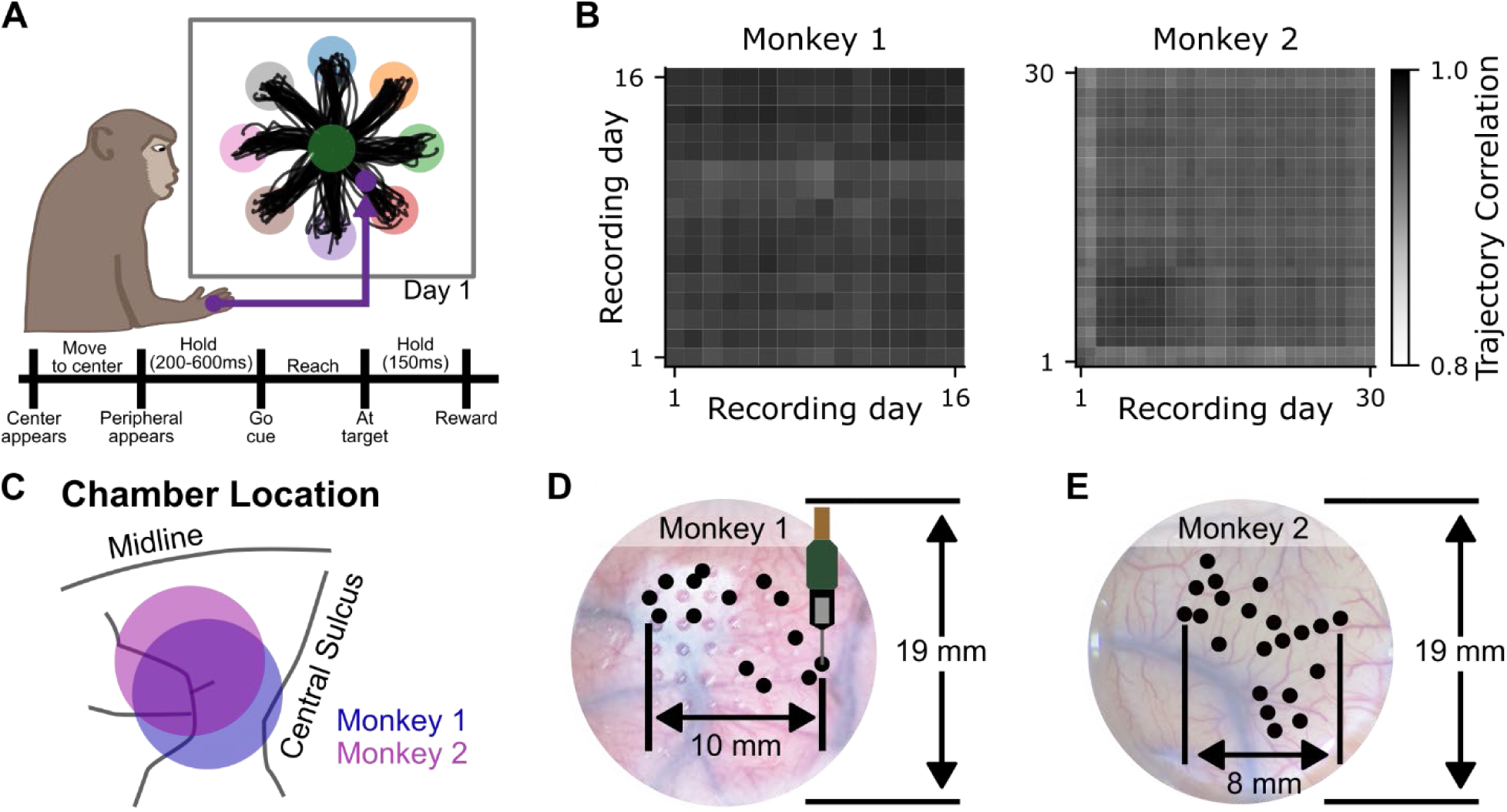
Large-scale recordings are made during highly stereotyped reaches. **(A)** Schematic of the experimental setup. Hand position was tracked and mapped to the position of the cursor on a visual display. The monkey performed a center-out reaching task with similar trajectories on trials to the same target (Monkey 1). Bottom: trial structure. **(B)** Average cursor trajectory correlation across days for each monkey. **(C)** Chamber location for each monkey. **(D, E)** Neuropixels probe insertion locations (black dots) on the cortical surface (recording sites) for Monkey 1 and Monkey 2, respectively.

### Behavioral recordings and task

Monkeys performed a self-paced delayed center-out reaching task with targets arranged in 2-dimensional plane (Fig. 1A). Movements of the monkey’s head and left arm were restricted during reaches. A glove with retroreflective markers was placed on the monkey’s right hand to allow its 3-dimensional position to be tracked using motion tracking cameras (NaturalPoint, Inc., Corvallis OR) and mapped to the position of a cursor on a screen (Ouchi et al., 2025). Upward hand movements moved the cursor towards the top of the screen and rightward hand movements moved the cursor towards the right of the screen. Trials began with the appearance of a center target. After a brief center hold, a peripheral target appeared 6.5 cm from the center target initiating the delay period. Subjects were required to keep the cursor in the center target during a delay period, the length of which was randomly drawn from a uniform distribution. Disappearance of the center target (the “go cue”) signaled animals to begin reaching to the peripheral target. A successful trial required a brief hold at the peripheral target, after which a juice or water reward was delivered. Peripheral targets were presented in the pseudo-randomized order to ensure an equal number of successful trials to each target. Task parameters used for each monkey can be found in Table 1.

**Table 1.** Center-out task parameters for each monkey.

| Parameter | Monkey 1 | Monkey 2 |
| --- | --- | --- |
| Target radius (cm) | 1.7 | 1.7-1.75 |
| Center target hold duration (ms) | 200 | 200 |
| Delay period range (ms) | 200 – 600 | 200 – 600 |
| Peripheral target hold duration (ms) | 175 | 100 |
| Reward duration (ms) | 100-200 | 200-250 |

### Behavioral data analysis

All monkeys were experts at this task during all recording sessions. Unrewarded trials and trials exceeding a maximum time threshold were removed from analysis (mean ± SD: 36.9 ± 19.2 (3.7 ± 1.9 %) removed trials for Monkey 1, 18.3 ± 18.6 (2.6 ± 1.9 %) removed trials for Monkey 2). The maximum trial length threshold was calculated to make the trial length distribution approximately symmetric with the following equation:

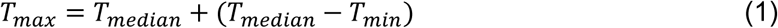

where *T* is the trial length, in seconds, between when the cursor enters the center target to initiate the trial to after the trial is rewarded. Median, minimum, and maximum trial length statistics were computed by pooling rewarded trials across all sessions.

We quantified stereotypy of performance across days by correlating the cursor trajectory from 100ms before to 400ms after the cursor left the center target. Data were aligned to movement onset time, approximated by the time when the cursor left the center target. Trials were separated by target direction, then we computed the Pearson correlation coefficient between each trial and all other trials to the same target, across all days. Pearson correlation was computed separately for the X and Y trajectory components, then averaged, weighted by the component variance:

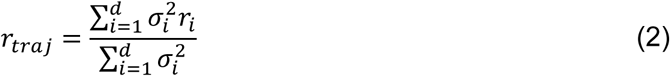

where *d* is the number of trajectory components, 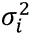 is the trajectory variance, and *r*_*i*_ is the Pearson correlation coefficient between two trajectories in the *i^th^* component.

### Electrophysiology

#### Neural recordings

We recorded neural activity using Neuropixels high-density laminar microelectrode arrays (IMEC, Leuven Belgium)(Jun et al., 2017). The Neuropixels probe was held by an oil hydraulic micromanipulator (Narishige International USA, Inc., Amityville, NY). A custom adaptor was designed in-house to securely mount the micromanipulator to the chamber and ensure the Neuropixels probe remained perpendicular to the implanted grid surface, which was approximately perpendicular to the cortical surface at the center of the chamber. We used the micromanipulator to manually translate the Neuropixels probe across the cortical surface until it was visually aligned with a selected grid hole. The probe was then slowly lowered with the micromanipulator to insert it into the brain.

We simultaneously recorded from 384 electrodes (of the 960 total electrode contacts on the Neuropixels probe) spanning 3.84 mm in depth during behavioral recordings. Our setup allowed us to insert approximately 4-5 mm (of the 10mm long Neuropixels probe) into the brain. We inserted probes and selected channels following procedures that aimed to align the top of the recorded electrode contacts with the cortical surface and to maximize the number of channels that recorded neural activity. We accomplished this in different ways for each animal. For Monkey 1, we recorded from the electrode contacts spanning the bottom 3.84 mm of the probe. We halted probe insertion once we recorded neurons within the top 0.2 mm of the recorded channels, even though the probe could have been physically advanced further. For Monkey 2, we inserted the probe as far as physically possible into the brain (approximately 4-5 mm), then took advantage of the ability to select the recorded electrodes online (in software). We based our channel selection on neural recordings from an initial baseline session where the 384 channels were configured to span 7.68 mm in depth. We used the rolling root-mean-square of the raw high-frequency (300-10,000 Hz; see below) signal computed by the data acquisition software (OpenEphys, Atlanta, Ga) across all electrodes, to estimate the location of the cortical surface using the empirical observation that electrodes not within the tissue would have clearly distinct statistics. Without moving the probe, we reconfigured the channel selection to select 384 channels spanning 3.84 mm of depth such that the most superficial neurons recorded during the baseline session were located within the top 0.2 mm of the recorded channels, thus aiming to match the configuration of Monkey 1. Baseline recordings were used offline to validate the online surface position estimation.

Electrical activity from each Neuropixels channel was hardware filtered into a high-frequency (AP) band (300-10,000 Hz) and sampled at 30 kHz (Jun et al., 2017). Electrical activity was then streamed to a dedicated computer via an external PXIe system (National Instruments, Austin, TX). Data acquisition was controlled on the dedicated neural acquisition computer which saved recorded activity locally (OpenEphys, Atlanta, GA).

We recorded from 14 and 21 unique grid locations, or recording sites, on the cortical surface for Monkey 1 and 2 respectively (Fig. 1D, E). We conducted these recordings on 16 and 30 non-consecutive days with multiple penetrations made at select grid locations. A majority of the data were collected using a single probe on each day except on four days with Monkey 2, where two probes were used to simultaneously record activity from different grid locations. Recording sessions were included in analyses using the following criteria: 1) they sampled neural activity from at least 2 mm along the Neuropixels probe, and 2) they contained at least 30 trials to each target that passed the trial length threshold. All analysis was conducted using 30 rewarded trials to each target.

#### Unit quality metrics

The raw AP band signals were post-processed to sort action potentials (spike sorting) using Kilosort 4.0 with default parameters (Pachitariu et al., 2024). Each putative neuron (unit) was then automatically assessed using quality metrics derived from the International Brain Laboratory (Table 2) (International Brain Laboratory, 2024). Units were only assessed during the trials selected for inclusion in the analysis. When there were more recorded trials than included in this analysis, we chose a start and end trial to qualitatively maximize stability. For most recordings, the chosen start trial was at the beginning of the behavioral session (median + SD: 0 + 0 trials for Monkey 1, 0 + 68 trials for Monkey 2). Units that passed all quality metrics were then manually inspected to ensure that the waveform did not clearly represent noise and that the firing rate did not contain sharp discontinuities across trials. Firing rate discontinuities were assumed to represent spike sorting errors and units exhibiting them were thus excluded. Only units that passed all quality metrics and manual inspection were used in further analysis.

**Table 2.** Unit quality metric description and acceptance thresholds.

| Quality Metric Name | Description | Acceptance Threshold |
| --- | --- | --- |
| Presence Ratio | The proportion of trials the neuron is active on. | = 1 (All trials included in analysis) |
| Refractory Period Violations | The number of spikes that violate the refractory period of the previous spike. | $\leq 1\%$ of spikes under 1ms |
| Waveform Amplitude | Peak-to-peak amplitude of the waveform on the channel that recorded the maximum amplitude. | $\geq 50 \mu\text{V}$ |
| Template Amplitude | The distribution of the waveform template amplitudes used by Kilosort. | Low bin threshold: $\leq 10\%$<br>Noise cut-off value: $\geq 5$ |

We computed the drift range (computed by Kilosort 4; (Pachitariu et al., 2024)) of each recording as a post-hoc proxy for recording stability and quality. Although Kilosort 4 corrects for drift, the drift range represents an estimation of Neuropixels probe movement relative to the brain and thus the recording instability.

We computed the signal-to-noise ratio (SNR) and true spike percentage for each unit as post-hoc proxies for sort quality. The SNR was defined as the ratio of the signal (average spike amplitude) over the “noise”. The recording noise was estimated by calculating the average median absolute deviation of the hardware filtered AP band data for channels at the same depth as the channel with the highest spike amplitude. The true spike percentage was defined as the percentage of spikes that were not refractory period violations (see *Table 2*).

#### Firing rate estimation

Firing rates of neurons were estimated for all units that passed all quality metrics by binning spikes into 10 ms bins. Firing rates for each unit were then z-scored across all time and trials, and smoothed using a Gaussian kernel with *σ* = 50 ms and a total kernel width of 3*σ*. This process used trial segments from 500 ms before to 1 s after the movement onset time.

#### Direction tuning analysis

We estimated how much each unit’s activity varied as a function of target direction by computing the modulation depth (MD) across different time periods within the trial. The MD was computed using the smoothed and normalized firing rate (150 ms moving window in 10 ms steps) and defined as the difference between the maximum and minimum firing rate across target directions.

### Decoding analyses

#### Linear decoding analysis

We quantified the task information represented in neural activity using target direction decoding analyses. This decoding analysis allowed us to identify units with low and high task information for visualizing relevant properties of the data and for future population dynamics analysis. Firing rates from 50 ms before to 250 ms after movement onset were averaged across time for each unit and each trial. This produced a matrix of neural activity with size (*n*_*Tr*_, *n*_*units*_), where *n*_*Tr*_ and *n*_*units*_ represent the number of trials and units, respectively. Linear decoding was performed on the time-averaged firing rates with linear discriminant analysis (LDA) implemented using the Python library Scikit-learn (Pedregosa et al., 2011) with the eigen solver and automatic shrinkage. Reported decoding accuracies were estimated by averaging the prediction accuracy from 4-fold cross validation. Each fold used 180 randomly selected trials to train the model and the remaining 60 trials to test the model. Chance estimation of decoding accuracy was computed using the same LDA decoding procedure applied to datasets with randomly shuffled target labels for each trial.

#### Nonlinear decoding analysis

Our primary results used linear decoding analyses. We supplemented this analysis with additional analyses using nonlinear decoding to quantify the task information as a control for the possibility that encoding nonlinearities may contribute to the observed spatial patterns of task information. Of the many possible non-linear decoding algorithms, we implemented the well-established latent factor analysis via dynamical system (LFADS) algorithm to classify target direction (Pandarinath et al., 2018). The LFADS model was implemented in TensorFlow and trained using an NVIDIA A40 GPU. We followed the same neural data preparation process as the linear decoding analysis (see *Electrophysiology*). To avoid the potential overfitting of nonlinear neural network models we implemented a 0.75/0.25 train/test split with dropout and L2 regularization. We additionally repeated model training and testing for 50 random initializations that randomly selected the units for analysis and randomly assigned trials to the train or test group. The reported decoding accuracy was estimated by averaging the accuracy of 50 random repeats. The total nonlinear analysis took approximately two days (one day for the dataset from each monkey) and the hyperparameter details are listed in Table 3.

**Table 3.** Nonlinear decoding model parameters.

| Parameter | Value | Description |
| --- | --- | --- |
| batch_size | 16 | Number of samples processed before model parameters update |
| num_layers | 2 | Number of hidden layers in the network |
| num_classes | 8 | Number of output targets for classification |
| num_epochs | 300 | Maximum number of training epochs |
| classifier_dims | 64, 32 | Dimensions of classifier layers |
| learning_rate | 1e-3 | Initial learning rate for optimization |
| hidden_dim | 64 | Dimension of hidden representations |
| n_factors | 16 | Number of factors |
| dropout_rate | 0.4 | Dropout probability for regularization |
| l2_reg | 1e-3 | L2 regularization coefficient |
| clip_value | 20.0 | Gradient clipping threshold |
| warmup_epochs | 10 | Number of warmup epochs for learning rate |
| min_warmup_lr | 1e-7 | Minimum learning rate during warmup |
| min_lr | 1e-7 | Minimum learning rate for scheduler |
| lr_patience | 50 | Learning rate scheduler epochs |
| lr_factor | 0.95 | Learning rate reduction factor |

#### Pseudo-population analyses

Our wide sampling across known locations on the cortical surface, paired with laminar recordings from the Neuropixels probes, provided 3-dimensional sampling across a large volume of the brain with precise known locations. We started with unit-level analyses by quantifying decoding accuracy for each unit separately. We then leveraged the animals’ consistent behavior across recording sessions to pool recorded units as pseudo-populations to quantify the spatial structure of task information and neural representations using decoding and population dynamics analyses (see below). To achieve our goals, we picked pseudo-population sizes that allowed us to assess the variation across neural populations. When feasible we looked at how trends were impacted by population size and saw that the relative trends persisted across a range of reasonable neural population sizes (Fig. S5E).

We analyzed the structure of task information across the cortical surface by quantifying target decoding accuracy of neural populations grouped by their cortical surface location. Cortical surface location (i.e., recording site) was defined using the grid location (spatial resolution 1 mm). To control for ensemble size, all populations included 25 units (Table 4). When more than 25 units were recorded at a given recording site, 25 units were randomly sampled and used for analysis to ensure decoding accuracy was estimated using the same number of units for each group; if fewer than 25 units were recorded at a recording site it was excluded from further analysis. Random unit sampling and analysis were repeated 1000 times, which was used to estimate the 95% confidence intervals.

**Table 4.**
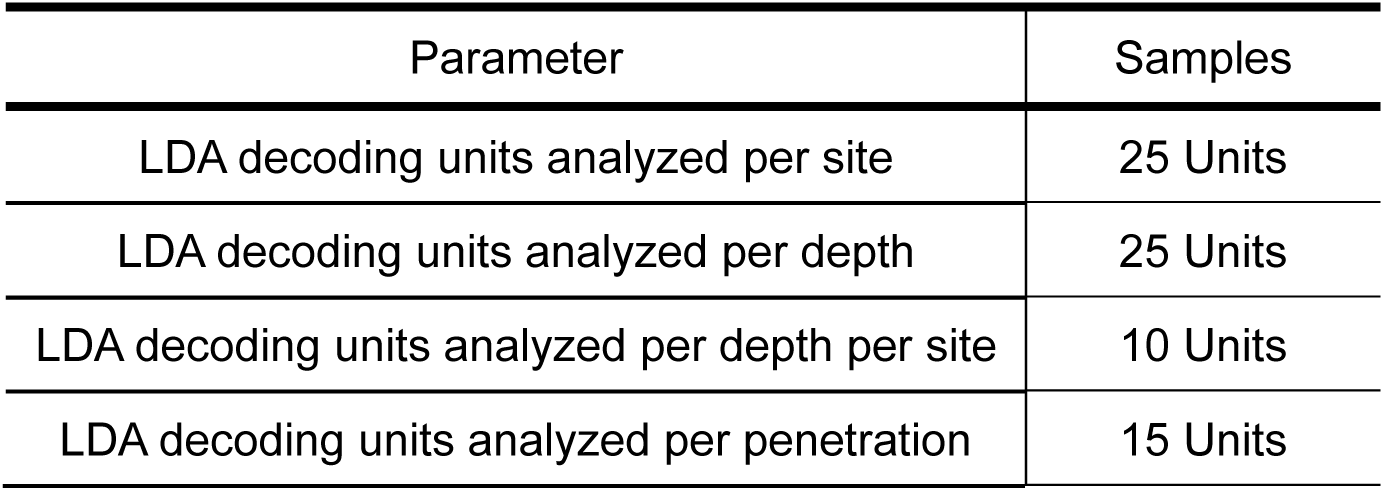
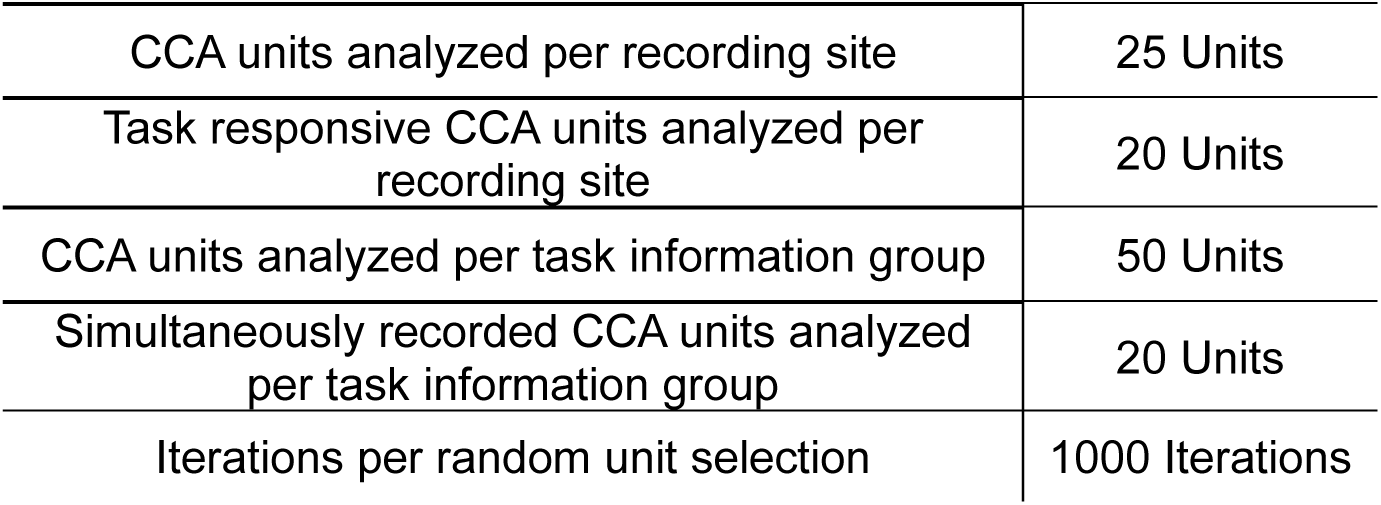
Analysis parameters sample size.

We controlled for firing rate differences across recording sites by sub-selecting the data to create similar firing rate distributions across recording sites. The distribution of the unit firing rates from each recording site was discretized with 10 Hz bins. The number of units selected from a bin for decoding accuracy calculations was set to the minimum number of units in each bin (*n*_*bin*_) across all sites. If a recording site had more than *n*_*bin*_ units for a given firing rate bin, *n*_*bin*_ units were randomly selected. Units were randomly selected and used to estimate decoding accuracy 1000 times to estimate a confidence interval for each recording site. This procedure led to decoding accuracy estimates that used neural ensembles of 16 units across all sites (both Monkey 1 and Monkey 2) with empirically-matched firing rates (R^2^: 0.96 for Monkey 1, 0.98 for Monkey 2; Fig. S2D).

We analyzed the structure of task information across cortical depths by quantifying target decoding accuracy of neural populations grouped by their location within the cortex relative to the estimated cortical surface. Units were selected using a similar strategy as described above to maintain a fixed ensemble size of 25 units (Table 4). Overlapping depth bins were constructed to span 1 mm with a 0.25 mm step size. Within each depth bin, units were randomly selected from across the cortical surface, creating a pseudo-population which was used to compute decoding accuracy. Unit selection and decoding analysis was repeated 1000 times, which was used to estimate the 95% confidence interval.

We also analyzed the structure of task information across both cortical surface location and depth. For these analyses, we repeated the analyses to compute decoding accuracy as a function of depth as described above but adjusted the unit ensemble size to 10 units given the reduced dataset size (Table 4). If fewer than 10 units were contained at any given depth, the 3D location was excluded from analyses.

We analyzed the variability in task information from repeated Neuropixels probe penetrations by quantifying target decoding accuracy with units grouped by the penetration. In each monkey, multiple penetrations were made at select recording sites. To control for ensemble size, all populations included 15 units (Table 4). When more than 15 units were recorded at a given recording site, 15 units were randomly sampled and used for analysis. When fewer than 15 units were recorded the recording was omitted from analysis.

### Neural population analysis

We applied canonical correlation analysis (CCA) to estimate the similarity of temporal dynamics between two neural populations, denoted as populations *A* and *B*. Neural populations *A* and *B* were created in a variety of ways (see below) and represent pseudo-populations in many cases. CCA was applied to Z-scored and smoothed firing rates (see *Firing Rate Estimation* for details) following procedures based on previous work (Gallego et al., 2018). Neural activity on individual trials between 50 ms before to 250 ms after movement onset were concatenated to create an array of size (*nt* ∗ *nTr*, *nunit*). *nt*, *nTr*, and *nunit* represent the numbers of time points within each trial, trials, and units in the neural population of interest, respectively. Principal Component Analysis (PCA) was applied to project the neural population firing rates onto the top 10 principal components (PC) and estimate the latent dynamics (*L*) for each neural population *A* and *B*, denoted as *L*_*A*_ and *L*_*B*_, respectively. The number of PCs was chosen based on prior literature (Gallego et al., 2020; Safaie et al., 2023), but we found that this hyper-parameter choice did not notably change our findings (Fig. S5A, B; S7A, B). We also controlled for explained variance by selecting the number of PCs for each neural population that accounted for 60% of the variance, which did not lead to notable changes to our findings (Fig. S5C, D).

CCA was then applied to “align” the latent dynamics of each neural population by finding the new directions that maximize the pairwise correlation between latent trajectories, referred to as the “aligned” axes (Gallego et al., 2018). CCA produces a correlation value for each mode that represents the similarity of the latent trajectories between each neural population. We report the CCA correlation using the average computed across the top 4 modes to minimize the possibility of spurious correlations (Sussillo et al., 2015; Gallego et al., 2020). We also used CCA to project the latent trajectories of each neural population pair (*L*_*A*_, *L*_*B*_) onto the “aligned” axes. After *L*_*A*_ and *L*_*B*_ were each projected onto the “aligned” axis, we then used LDA decoders as described previously (see *Decoding analysis*) to estimate the “aligned decoding accuracy”. We trained the decoder using all data from *L*_*A*_ and tested it on all data from *L*_*B*_ to assess decoder generalization to unseen data from a different neural population. The aligned decoding accuracy therefore quantifies the accuracy of a decoder trained on aligned neural activity from neural population *A* when tested on the aligned neural activity from neural population *B* for prediction.

#### Shared neural population dynamics between recording sites

We analyzed the spatial structure of population dynamics by performing CCA alignment analyses on different pseudo-populations following a similar approach as the linear decoding analyses (see *Psuedo-population analyses*). Pseudo-populations with a fixed ensemble size were created by randomly selecting 25 units from each recording site (Table 4). We used the randomly selected units to perform CCA between the neural populations recorded at each pair of recording sites. Each reported CCA correlation quantifies the similarity of the temporal dynamics across the neural populations. We repeated this analysis 1,000 times and randomly selected different units on each iteration to estimate the 95% confidence interval. When analysis was restricted to task-responsive units we used 20 units from each recording site to ensure consistent population sizes.

We performed three comparisons to assess patterns in CCA correlation between neural populations from different recording sites. First, we compared the absolute distance between recording site grid locations to the CCA correlation of each pair of neural populations (Fig. 3C, D). Second, we organized neural populations by their rostral-caudal axis location (Fig 3.E, F). To assess how the rostral-caudal location of an individual neural population is related to CCA correlation, we compared the CCA correlation between that neural population and its neighbors: the next most rostral neural population, and the next most caudal neural population. We averaged the CCA correlation between each pair of these three neural populations and report that as the Neighboring CCA correlation based on rostral-caudal position (Fig. 3G). Third, we organized neural populations by their decoding accuracy (see *Linear decoding analysis*; Fig. 3H, I). To assess how the task information of an individual neural population is related to CCA correlation, we compared the CCA correlation between that neural population and its neighbors: the neural population with the next lowest decoding accuracy and the neural population with the next highest decoding accuracy (see Fig. 2G, H). We averaged the CCA correlation between each pair of these three neural populations and report that as the Neighboring CCA correlation based on decoding accuracy (Fig. 3J).

**Figure 2.**
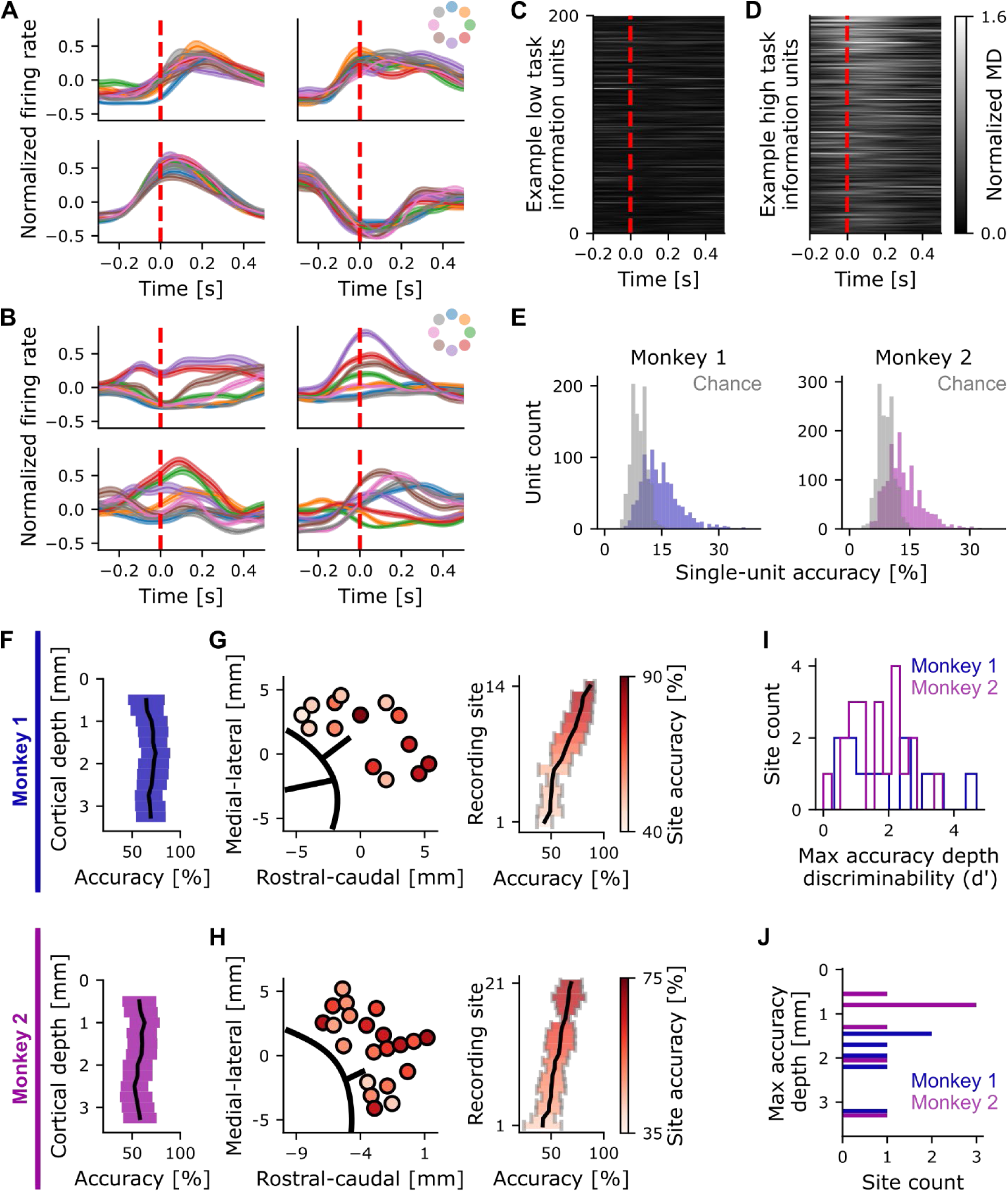
Task information is heterogeneously distributed across frontal motor cortices. **(A)** The average response of four example units at movement onset of reaches towards each target with qualitatively little separation in the response across different targets (low task information). Each color represents the trial-averaged neural activity to different targets. Red dashes indicate the time of movement onset. Inset in the upper right represents the target that each color is assigned to. Solid lines indicate the mean with shading indicating the standard error of the mean across trials. **(B)** Same as (A) but for different example units with qualitatively large separation in response to reaches towards different targets (high task information, see *Linear decoding analysis*). **(C)** Modulation depth (MD) (calculated using normalized firing rates) as a function of time for 200 units whose activity was the least predictive of the reach target direction (see *Linear decoding analysis*) during movement. Firing rates were time-aligned to movement onset, indicated by the red dashed line. **(D)** Same as (C) but for a different population of 200 example units that exhibited high task information by best predicting reach target direction. **(E)** Distribution of single-unit decoding accuracy across all recorded units for each monkey (monkey 1, left, blue; monkey 2, right, pink) overlayed with the chance decoding distribution (grey). **(F)** Distribution of population decoding accuracy across depths. Units at the same depth (+/- 500 µm) across recordings were compiled into distinct neural populations from which decoding accuracy was estimated. Black lines indicate the mean; the shaded region represents the 95% confidence interval (CI). **(G)** Left: Distribution of population decoding accuracy across the cortical surface (site accuracy) for monkey 1. Units from the same recording site were compiled into distinct neural populations from which decoding accuracy was estimated. The black lines represent the arcuate sulcus. Right: Population decoding accuracy across recording sites, sorted by the mean decoding accuracy. Black lines indicate the mean; the shaded region represents the 95% CI. Colors indicate the site decoding accuracy (color bar) for both right and left panels. **(H)** Same as (G) for monkey 2. **(I)** Distribution of discriminability index (d’) between the depth range with the maximum decoding accuracy and the remaining units (see *Statistical significance estimation*) across recording sites (N=12 for monkey 1; N=21 for monkey 2). **(J)** Histogram of the depth (+/- 500 µm) where the highest decoding accuracy was observed for recording sites where decoding depth significantly varied across depths (N=6 for monkey 1; N=7 for monkey 2).

**Figure 3.**
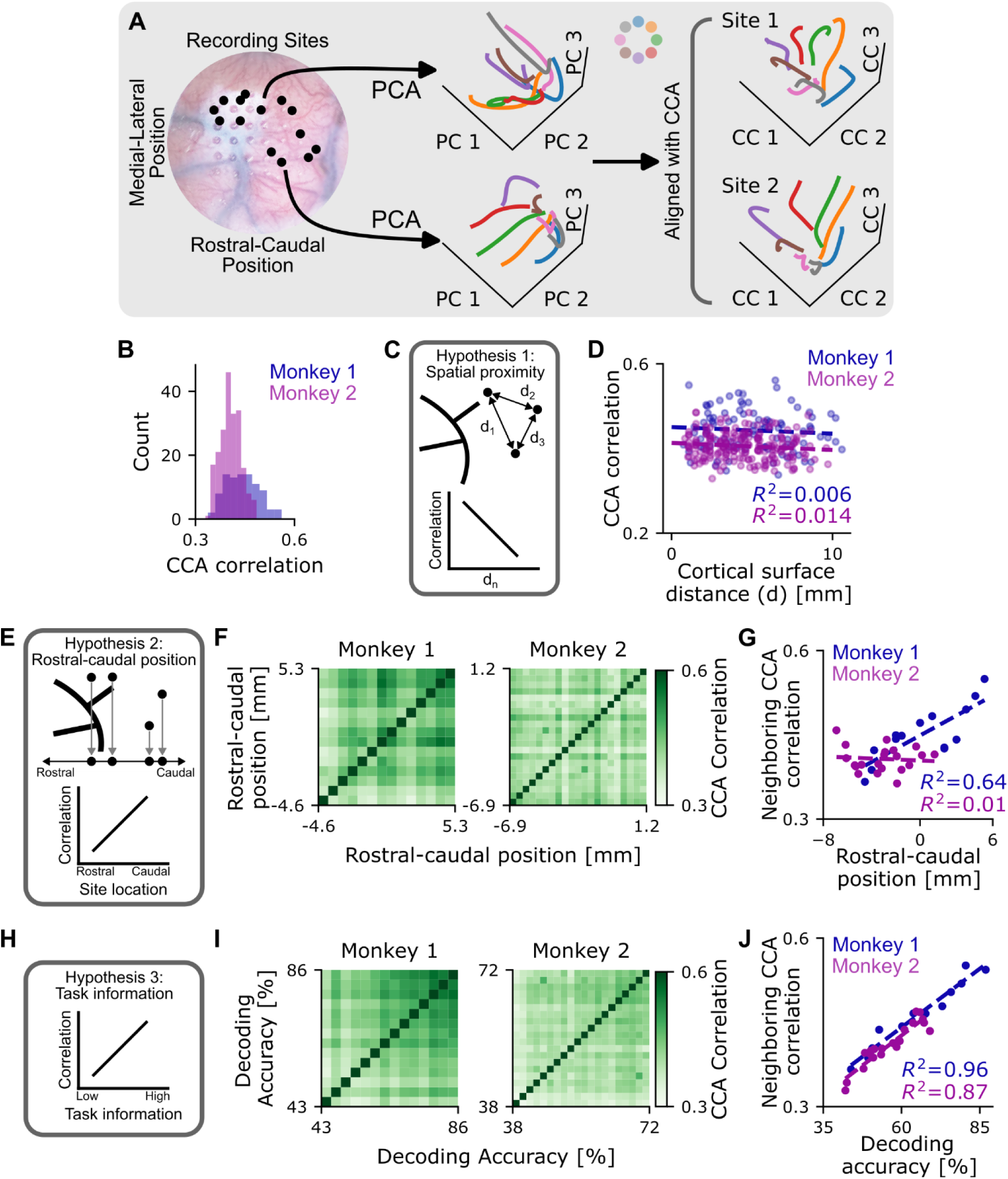
Task information predicts similar dynamics better than spatial position. **(A)** Diagram of canonical correlation analysis (CCA) of latent dynamics to quantify their similarity. Recording site locations for Monkey 1 (left), the latent dynamics of distinct neural populations each composed of units recorded at different recording sites (middle), and the latent dynamics of each neural population projected onto the shared axes found with CCA (right). Each color represents the trial-averaged dynamics to a different target (inset). **(B)** The distribution of CCA correlation between pairs of neural populations from different recording sites. **(C)** Diagram depicting hypothesis 1. Each dot (top) represents a recording site and d_n_ represents the absolute distance between a pair of recording sites. This hypothesis predicts that the CCA correlation decreases as two neural populations become further apart (bottom). **(D)** Relationship between CCA correlation and the spatial proximity of two distinct neural populations for monkey 1 (blue) and monkey 2 (pink). Each dot represents a pair of neural populations (N=91 for monkey 1; N=210 for monkey 2); dashed lines represent the linear regression. **(E)** Diagram depicting hypothesis 2, formatted as in (C). This hypothesis predicts that CCA correlation increases as location moves caudally along the rostral-caudal axis. **(F)** Heatmap of CCA correlation between neural populations organized by rostral-caudal position for Monkey 1 (left) and Monkey 2 (right). Each pixel represents a pair of neural populations (N=91 for monkey 1; N=210 for monkey 2). **(G)** Neighboring CCA correlation based on rostral-caudal position for each distinct neural population (see *Neural population dynamics between recording sites*) for monkey 1 (blue) and monkey 2 (pink). Each dot represents a recording site (N=14 for monkey 1; N=21 for monkey 2). **(H)** Diagram depicting hypothesis 3, formatted as in (C). This hypothesis predicts that CCA correlation increases as the amount of information contained within a population increases. **(I)** Heatmap of CCA correlation between distinct neural populations organized by the amount of task information for Monkey 1 (left) and Monkey 2 (right). Format as in (F). **(J)** Neighboring CCA correlation based on task information for each distinct neural population, formatted as in (G).

As an additional control, we generalized the rostral-caudal CCA analyses above to look for structure along any axis on the cortical surface. We first defined 1,000 random axes on the cortical surface, then we projected the spatial position of each neural population onto each random axis to get their relative location on the new, random axis. We then regressed the neighboring CCA correlations with the distance along the (randomly chosen) spatial axis and computed the coefficient of determination (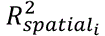).

We compared whether decoding or spatial organization better predicts the neural populations with similar dynamics by computing the Task Index for each spatial axis:

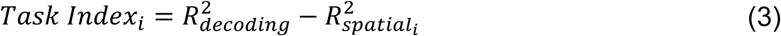

where 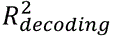 is the coefficient of determination of CCA with difference in task decoding. Positive *Task Index* values suggest that similar dynamics are best predicted by decoding accuracy while negative values suggest that similar dynamics are best predicted by spatial organization.

#### Shared neural population dynamics between task-tuned populations

We assessed the temporal dynamics of task-tuned units by creating neural pseudo-populations comprised of units with either low or high task information. We used the single-unit decoding accuracy (see *Linear decoding analysis*) to define the amount of task information encoded by each recorded unit. The 100 units with the highest single-unit decoding accuracy were defined as the *unit-level high task information population*. The 100 units with the lowest single-unit decoding accuracy were correspondingly defined as the *unit-level low task information population*. These units were then arbitrarily split into two groups with low task information (50 units each) and two groups with high task information (50 units each) for further analysis (Table 4). We computed the CCA correlation (Fig. 4B, C) and aligned decoding performance between each combination of these unit groups (i.e., low-low, high-high, low-high; Fig. 5). Random unit splitting and analysis was repeated 1000 times to estimate a 95% confidence interval. We assessed the statistical significance of dynamic correlation estimates using CCA lower bounds (see *CCA lower bound estimation*).

**Figure 4.**
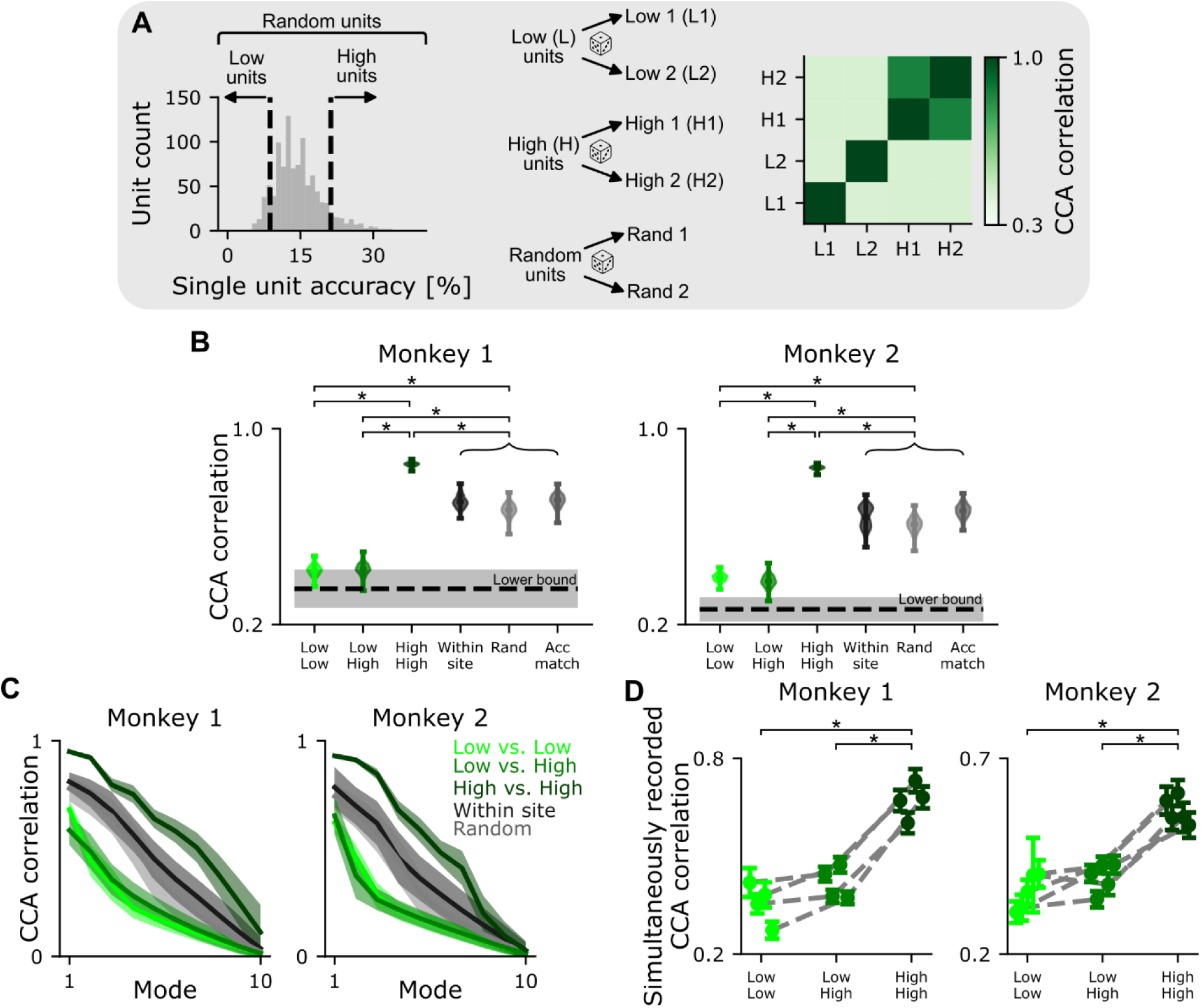
Neural populations composed of units with high task information have similar dynamics. **(A)** Diagram depicting how distinct neural populations were composed and compared. Units were grouped based on their single-unit decoding accuracy into high (H) and low (L) task information groups (left). Each of these groups were randomly split into distinct neural populations as an internal control (L1, L2, H1, H2). We then compared the dynamics between each combination of these four distinct neural populations using CCA for all population combinations defined by task information (right). Neural populations composed of units drawn from all recorded units were also composed and compared (Rand). **(B)** CCA correlation between distinct neural populations defined by task information (L and H), recording site (Within site), random selection (Rand), or random selection while matching population decoding accuracy to the high task information populations (Acc match) for monkey 1 (left) and monkey 2 (right). The distribution for each comparison represents the variability from different random unit selections (N=1000). Each distribution depicts the CCA correlation results from all random unit selections and is centered on the mean. The black dashed line represents the lower bound estimated from inter-trial interval activity (see *CCA lower bound estimation*) and the gray shading represents the 95% CI across random unit selections. **(C)** CCA correlation as a function of CC mode for select neural population comparisons. The shaded regions represent the 95% confidence interval for each mode across random unit selections. **(D)** CCA correlation restricted to populations of simultaneously recorded units. Units were split based on their task information into new high and low task information groups for comparison (see *Shared neural population dynamics between task-tuned populations*). The dashed line connects comparisons from the same recording site and error bars represent the 95% CI across random unit selections. Significance labels represent when all distributions, matched for recording site, were significantly different.

**Figure 5.**
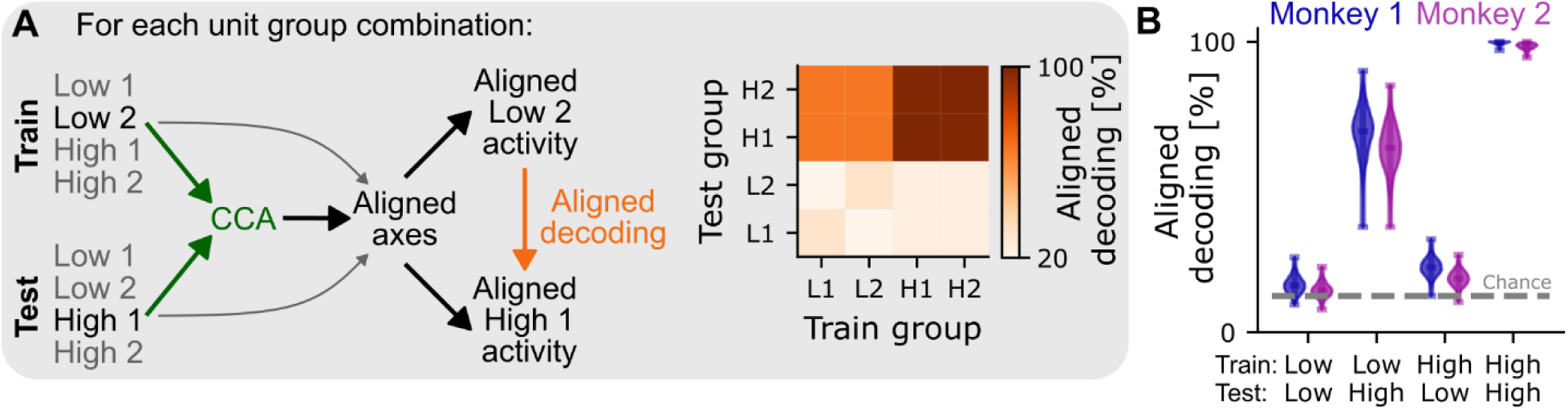
Neural populations composed of spatially distributed units with high task information have similar task representations. **(A)** Diagram depicting analyses to assess whether neural populations have similar task representations in their latent dynamics (left). We applied CCA to the neural populations composed of units with either high or low task information to find the axes that best align the individual latent dynamics (“aligned axes”). We then projected the latent dynamics from each neural population onto the aligned axes (gray arrow), trained a decoder using aligned activity from one population, then tested the resulting decoder using aligned activity from the other population (“aligned decoding”). Heatmap representation of the resulting aligned decoding between all combinations of neural populations (right). **(B)** Aligned decoding for each neural population comparison. The distribution represents the variability of aligned decoding from different random unit selection (N=1000). Each distribution depicts the aligned decoding results from all random unit selections and is centered on the mean. The gray dashed line represents chance-level decoding.

To ensure our results were not driven by how units were assigned to populations (i.e. high task information or low task information populations), we applied CCA to neural populations composed of random unit groups, defined as the *random unit population*. Two distinct neural populations were constructed by randomly selecting recorded units without replacement. We repeated random unit selection and CCA analysis 1000 times to estimate the 95% confidence interval (Fig. 4B, C).

We estimated the site-level dynamics similarity by performing CCA on neural populations composed of units recorded from the same site. We constructed two distinct populations of units recorded at the same site by randomly selecting units without replacement. We repeated random unit selection and CCA analysis 1000 times to estimate the 95% confidence interval (Fig. 4B, C).

We also replicated our CCA analyses restricting our data to simultaneously recorded neural populations to assess whether pseudo-population analyses underestimated or missed shared dynamics (e.g., due to non-task related neural activity that varies more from session to session). We restricted our analysis to simultaneously recorded units, then split units into high and low task information groups using the procedure described above. We used populations of 20 units to control for population size and recordings with fewer than 80 simultaneously recorded units were excluded from analysis. Random unit selection and CAA analysis was repeated 1000 times to estimate a 95% confidence interval.

### Statistical analysis

#### CCA lower bound estimation

We sub-selected data from timepoints outside the task to assess the specificity of correlated dynamics between two neural populations to the task (Safaie et al., 2023). We used random timepoints selected from inter-trial interval data, which was defined as the time interval between the end of the previous trial and when the center target appeared to initiate the next trial session (mean ± SD: 0.41 ± 0.88 s for Monkey 1, 0.39 ± 0.72 s for Monkey 2). The neural activity of randomly selected units during these task intervals was concatenated and analyzed using the same procedure as if aligned to behavior (see *Neural population analysis*). Units were randomly chosen 1000 times to estimate the 95% confidence interval of CCA correlation.

#### Statistical significance estimation

We used bootstrapping to estimate the underlying statistical distributions of decoding, CCA correlation, and aligned decoding metrics. We considered two distributions significantly different if their 95% confidence intervals had zero overlap. Analyses results were considered not statistically different from chance levels if the 95% confidence interval included chance performance. Chance performance for a classifier applied to neural populations was assumed to be equivalent to random target selection (12.5% decoding accuracy for an 8-target task with approximately uniform target presentation).

We quantified the difference in decoding accuracy between depths at each recording site using the discriminability index (d’) (Williams et al., 2013). The d’ metric quantifies the separation between two statistical distributions (groups 1 and 2) with means *μ*_1_, *μ*_2_ and standard deviations *σ*_1_, *σ*_2_ via:

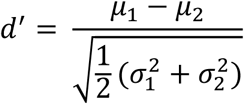

The d’ value represents how many standard deviations apart the means of the two distributions are, weighted for unequal variances. We used bootstrap resampling to estimate the underlying statistical distribution of decoding accuracy for two groups at each recording site: 1. units in the depth range with the highest decoding accuracy (see *Pseudo-population analysis*) and 2. all remaining units. We then computed the d’ of these two distributions (Fig. 2I). We also used bootstrap resampling to estimate a chance distribution of d’ from each recording site. Units were randomly assigned to two groups (A, B), mirroring the ‘high decoding depth’ and ‘remaining units’ groups described above. Random unit assignment and the chance d’ calculation were repeated 1000 times for each recording site. We tested if the empirical d’ was significantly greater than chance by computing the one-sided p-value (alpha = 0.05) separately for each site. A significant p-value would therefore suggest that units in the ‘high decoding depth’ had significantly higher decoding accuracy than the remaining units. Recording sites with less than 10 units outside the chosen depth were excluded from this analysis.

#### Statistical analysis across multiple penetrations

We assessed the potential variability in our results that could be attributed to the random sampling of neurons at a given recording site inherent in electrophysiology by analyzing data collected from the same site across different sessions (i.e., repeated insertions of the Neuropixels probe into the same approximate location on different days). We compared the decoding accuracy difference between multiple penetrations within the same recording site to the decoding accuracy difference between penetrations at other recording sites (Fig. S3D, E). We estimated the decoding accuracy difference for each pair of repeated penetrations at the same recording site with

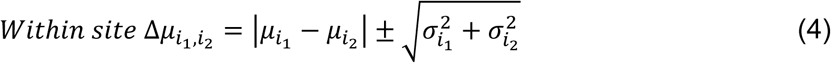

where *μ*_*i*_1__, *μ*_*i*_2__ and *σ*_*i*_1__ *σ*_*i*_2__ represent the mean decoding accuracy and standard deviation, respectively, across unit resampling for two separate penetrations at recording site *i*_*j*_.

We estimated the decoding accuracy difference across penetrations from different recording sites for penetration *i*_1_ by computing the decoding accuracy difference between penetration *i*_1_ and all other penetrations performed at different recording sites.

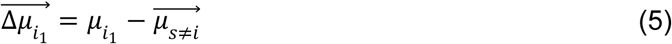

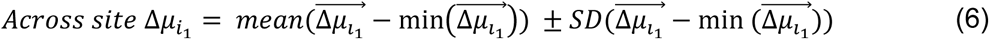

In equation 5, *μ*_*i*_1__ represents the mean decoding accuracy from penetration 1 at recording site *i* across unit resampling, and 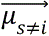 represents the vector of mean decoding accuracy across unit resampling from all penetrations at all recording sites, *s*, except when conducted at the same recording site *i*. We used equation 5 to compute the reported across-site decoding accuracy difference and standard deviation through equation 6, where *mean* and *SD* indicate that the mean and the standard deviation were computed across all recording sites, *s*.

#### Task-responsive unit estimation

We estimated if individual units were modulated by the task (i.e. task responsive) by comparing the activity around movement onset with baseline activity on a single trial basis using a signal detection theory approach (Banerjee et al., 2010). Briefly, this approach computes the cumulative time-varying log-likelihood ratio (AccLLR) to test whether the single-trial firing rate (before normalization and smoothing) better fits the mean baseline activity or the mean movement onset activity. The activity from 500 ms to 100 ms before movement onset was considered the baseline window. The activity from movement onset to 400 ms after movement onset was considered the response window. We assessed whether a unit had a significant response on a trial by trial basis by comparing the AccLLR during the response window to a detection threshold calculated using receiver operator characteristic (ROC) analysis, which compared the probability of correct to incorrect response detection. Units were considered task-responsive if the area under the curve (AUC) from the ROC analysis was significantly greater than the null hypothesis of AUC = 0.5 using a one-sided Z-test with false discovery rate correction (p ≤ 0.05).

## Results

We trained two monkeys to perform a center-out reaching task (Fig. 1A). Both monkeys were proficient in the task and produced highly stereotyped movements (reach time mean ± SD: 0.19 ± 0.07 s for Monkey 1, and 0.18 ± 0.07 s for Monkey 2). The average pairwise correlation between the individual trial movement trajectories across all trials and all days was near 1 for both the 2D cursor position (mean ± SD: 0.96 ± 0.04 for Monkey 1, 0.94 ± 0.07 for Monkey 2; Fig. 1B, S1A) and 3D hand position (mean ± SD: 0.96 ± 0.05 for Monkey 1, 0.94 ± 0.07 for Monkey 2; Fig. S1B), confirming consistent reach trajectories across all recording days.

We recorded single-unit extracellular activity from chambers implanted over frontal motor cortices while the monkeys performed center-out reaches (Fig. 1C; see *Behavioral recordings and task*). We used high-density laminar microelectrode arrays (Neuropixels) to make spatially precise recordings of neural populations from a large 3D cortical volume (Fig. 1D). Neural recordings covered a cortical surface area of 9.9 × 6.7 mm in Monkey 1 and 8.1 × 9.3 mm in Monkey 2 (rostral–caudal × medial–lateral; Fig. 1D, E). Each probe penetration recorded activity from the cortical surface to a depth ranging from 2.0 - 3.8 mm. Collectively, we recorded 1,022 units from Monkey 1 and 1,441 units from Monkey 2. The behavior stereotypy, spatially precise recordings across a broad cortical region, and large quantity of units allowed us to group neurons into pseudo-populations, then compare the spatial organization of task related information and the temporal structure of neural dynamics across frontal motor cortex.

### Task information is heterogeneously distributed across frontal motor cortices

We first examined how much individual units modulated their activity with reach direction to assess the neural encoding of one key component of task information. Trial-averaged and target-specific peri-stimulus time histograms revealed clear differences in target direction modulation across units: some units showed task modulation, but qualitatively little difference in neural activity between targets (Fig. 2A), while other units had qualitatively distinct responses during reaches to different targets (Fig. 2B). We computed the modulation depth (MD) for each unit across time within the trial (see *Direction tuning analysis*) and found that these example units were representative of broader differences in target direction modulation across hundreds of other units (Fig. 2C, D). This qualitative evidence suggests that variability in the degree to which individual units modulated their activity with target direction, which we will refer to as task information, was present across all recorded units.

We quantified the distribution of task information across all recorded units by computing how well each individual unit predicted target direction (decoding accuracy; see *Linear decoding analysis*). The distribution of single-unit decoding accuracy was non-normal (Kolmogorov-Smirnov test: p < 0.0001 for Monkey 1, p < 0.0001 for Monkey 2) and showed a statistically distinct rightward skew compared to the chance distribution (Skewness ± S.E.: 1.17 ± 0.08 vs. chance: 0.19 ± 0.08 for Monkey 1; 1.19 ± 0.06 vs. chance: 0.25 ± 0.06 for Monkey 2, Fig. 2E). Single-unit decoding accuracy was not strongly correlated with the average unit firing rate (R^2^: 0.07 for Monkey 1, and 0.07 for Monkey 2; Fig. S2A) or proxies for spike-sorting quality like signal-to-noise ratio (SNR; Fig. S3A) and the percentage of true spikes based on refractory period violations (Fig. S3B). These results align with previous findings that many units in motor cortex are untuned (Georgopoulos et al., 1982; Best et al., 2016). We next leveraged our high density, spatially-precise recordings to investigate the spatial organization of task information across neural populations while controlling for population size.

We grouped units into neural populations based on the depth they were recorded from, irrespective of cortical surface location (i.e., recording site). Our dense sampling found that no single depth consistently encoded more task information than others (decoding accuracy confidence intervals (CI) overlapped across depths; see *Statistical significance estimation*; Fig. 2F), in contrast to previous reports suggesting depth-dependent patterns among more sparsely sampled neurons in motor cortices (Mollazadeh et al., 2011). We then regrouped units into neural populations defined by the recording site, independent of depth. Although all recording sites showed decoding accuracy above the 12.5% chance level, some locations had significantly higher decoding performance than others (no CI overlap across locations; see *Statistical significance estimation*; Fig. 2G, H). To rule out the possibility that these differences were due to our methods, which assume linear encoding, we repeated this analysis using a nonlinear neural network (see *Nonlinear decoding analysis*; Fig. S4). We found a similar range of population decoding accuracy across sites with this alternate decoding approach. Additional controls confirmed that task information decoding was not strongly correlated with the total number of units (R^2^: 0.20 for Monkey 1, and 0.01 for Monkey 2; Fig. S2B), the mean firing rate (R^2^: 0.37 for Monkey 1, and 0.01 for Monkey 2; Fig. S2C-G), or proxies for recording quality like the recording drift range estimated by Kilosort 4 (R^2^: 0.05 for Monkey 1, and 0.14 for Monkey 2; Fig. S3C), suggesting the decoding accuracy differences were primarily driven by recording site. Moreover, the decoding accuracy variability across recording sites was larger than that for repeated penetrations at the same recording site, for each of 5 recording sites with multiple penetrations (Fig S3D, E).

Based on the variability in task information across recording sites, we revisited the influence of cortical depth, now analyzing depth-dependence individually for each recording site. We identified the 1 mm depth range with the highest decoding accuracy for each site, which we compared to the decoding accuracy obtained with all other units using the discriminability index (d’) (see *Statistical significance estimation*). This metric captures the separation between these two distributions (in terms of standard deviations, weighted for unequal variance). We observed a range of discriminability across recording sites, suggesting that many sites had a depth with higher decoding than the remaining units (Fig. 2I). We tested whether this depth-dependence was statistically significant by comparing the observed d’ to a chance distribution (see *Statistical significance estimation*). We found that some sites had minimal depth-dependence (N = 6 of 12 for Monkey 1, and N = 14 of 21 for Monkey 2). For sites with depth-dependence (N = 6 of 12 for Monkey 1, and N = 7 of 21 for Monkey 2), the highest decoding accuracy spanned multiple millimeters for each Monkey (Fig. 2J). Our observations reveal that the depth-decoding accuracy relationship varied across recording sites. Collectively, our results suggest that task information was heterogeneously distributed across both the motor cortical surface and in depth.

### Task information predicts similar dynamics better than spatial position

The prior analyses suggest that task information was unevenly distributed at the level of single neurons. Temporal patterns of motor cortex activity during movement production similarly appeared unstructured at the level of single neurons (Churchland and Shenoy, 2007), yet have been shown to underly more coordinated dynamics at the level of neural populations (Churchland et al., 2012; Kaufman et al., 2014; Elsayed et al., 2016; Lara et al., 2018). We therefore asked whether coordination at the level of population dynamics may shed light on the spatial heterogeneity of task information in individual neurons and neural populations.

We first compared population temporal dynamics across different neural populations, grouping neurons based on their recording site, which partially captured the spatial variation in task information at the single-unit level (see Fig. 2). We assessed the similarity of population dynamics using canonical correlation analysis (CCA) (see *Neural population analysis*; Fig. 3A). We estimated each neural population’s latent dynamics and then compared the latent trajectories across each pair of neural populations with CCA. We found that the similarity of population dynamics notably varied across population pairs grouped by recording site (mean ± SD: 0.44 ± 0.05 for Monkey 1, 0.41 ± 0.03 for Monkey 2; Fig. 3B). Estimated CIs on population similarity showed no overlap between the most (95% CI: [0.51, 0.61] for Monkey 1, [0.43, 0.54] for Monkey 2) and least (95% CI: [0.30, 0.38] for Monkey 1, [0.28, 0.38] for Monkey 2) correlated neural populations across the datasets, suggesting statistically meaningful differences. We note that our reported CCA values are lower than common reports in motor areas (Gallego et al., 2020; Safaie et al., 2023) due to our controls for neural population size, which influences CCA correlations (Fig. S5E).

We investigated three hypotheses to identify structure that might explain these seemingly heterogeneous temporal dynamics. First, we hypothesized that dynamics similarity was related to spatial proximity of neural populations (Fig. 3C), motivated by past observations that nearby neurons are likely to be anatomically connected (Gatter and Powell, 1978) and tend to have similar task information encoding properties (e.g., preferred directions (Amirikian and Georgopoulos, 2003)). We compared the pairwise CCA correlation of two neural populations to the absolute distance across the cortical surface separating those populations, which revealed no clear trends (R^2^: 0.006 for Monkey 1, 0.014 for Monkey 2; Fig. 3D). Both near and distant neural populations had correlated dynamics, suggesting spatial proximity alone does not explain correlated dynamics.

Second, we hypothesized that dynamics similarity was related to the location of the neural populations along an anatomically-defined axis across the cortical surface (Fig. 3E). We focused on the rostral-caudal axis based on the anatomical and functional organization of brain regions in frontal motor cortices (Gatter and Powell, 1978; Riehle and Requin, 1989; Dum and Strick, 2002; Cisek and Kalaska, 2005). We found that neural populations organized by their location along the rostral-caudal axis showed some relationship to dynamics similarity, but the relationships were not consistent across monkeys. Caudal neural populations had qualitatively more similar dynamics than rostral neural populations, consistent with our hypothesis, in Monkey 1. However, this pattern was not observed in Monkey 2. (Fig. 3F). We quantified this pattern by computing the Neighboring CCA correlation, which computes how correlated the dynamics of a given neural population are to neural populations with nearby rostral-caudal positions (see *Shared neural population dynamics between recording sites*; Fig. 3G). This revealed a clear correlation between rostral-caudal location and population dynamics in Monkey 1 (R^2^ = 0.64), but no trend in Monkey 2 (R^2^ = 0.01), potentially related to slightly different chamber locations (see Fig. 1C). These results suggest that the rostral-caudal location of a neural population likely impacts how similar the dynamics of two neural populations are, yet the inconsistent results across monkeys suggest other factors are involved.

Third, we hypothesized that dynamics similarity was related to task information encoding within the neural population (Fig. 3H). This hypothesis is consistent with populations of neurons coordinating to perform functional computations, potentially independent of spatial location. Such an architecture is supported by past observations of mixtures of spatially local and distributed task information in multi-neuron resolution functional mapping studies (Chehade and Gharbawie, 2023; Ouchi et al., 2025) and anatomical connections spanning multiple millimeters that could support spatially-distributed coordination (Gatter and Powell, 1978; Huntley and Jones, 1991). We organized neural populations by their task information and observed that neural populations with high task information had correlated dynamics while neural populations with low task information tended to have distinct dynamics (Fig. 3I). We quantified this pattern by computing the Neighboring CCA correlation with neural populations organized by task information, rather than spatial position, and saw a strong trend in both animals (R^2^ = 0.96 for Monkey 1, R^2^ = 0.87 for Monkey 2). This analysis highlights that spatially distributed neural populations had correlated dynamics if they were task modulated, suggesting that the temporal dynamics between neural populations is best predicted by the task information of the neural populations rather than spatial location. To ensure these results were not specific to the particular coordinate system we chose (i.e., anatomical axis), we repeated the spatial analysis using different axes across the cortical surface. We found that task information was always a better predictor of dynamics similarity than any axis across the cortical surface (Fig. S6C, D).

Collectively, our results revealed that the dynamics of neural populations varied, but similar dynamics were best predicted by the underlying variation in task information that they contained. Distinct neural populations had correlated dynamics if they had similarly high amounts of task information, even if the neural populations were spatially separated. We found qualitatively similar findings when replicating these analyses with neural populations composed of only task-responsive units (see *Task-responsive unit estimation*), suggesting that task-responsiveness does not explain the patterns of dynamics similarity (Fig. S7D-G). The relationship between dynamics similarity and task information we observed also does not appear to reflect a trivial correlation between these potentially-related metrics. Conceptually, correlated dynamics and task information can be independent (Fig. S8A-C). Empirically, we observed only weak correlations between task information and dynamics similarity when generating neural populations via random selection (R^2^: 0.14 for Monkey 1, and 0.15 for Monkey 2; Fig. S8D, E).

### Similar dynamics are driven by a subset of units with high task information

Our results suggest that task information played a role in producing correlated dynamics in distinct neural populations. However, there are several possible explanations for the correlation we observed between the site-level task information and population dynamics. For instance, this link could be at the unit level, where a select few highly task modulated units drove the coordinated dynamics, or at the site level, where all units at a recording site collectively contributed to coordinated dynamics. To distinguish these alternatives, we pooled all recorded units and then separated them only by their task information, which was estimated using single-unit decoding accuracy (see *Shared neural population dynamics between task-tuned* populations; Fig. 4A). We created two pseudo-populations with high and low task information by grouping together the 100 units with the highest and lowest single-unit decoding accuracy, respectively. As an internal control, we further separated these unit-level high and low task information neural populations into two distinct populations (random assignment) to create two high and two low neural populations (four total). We analyzed how task information influenced neural population dynamics by comparing each neural population (Fig. 4A) while controlling for the number of units in each neural population (Fig. S5E).

We found that unit-level high task information populations had correlated dynamics (95% CI: [0.84, 0.87] for Monkey 1, [0.83, 0.85] for Monkey 2; High vs High in Fig. 4B). Unit-level low task information populations, in contrast, exhibited more distinct dynamics (95% CI: [0.38, 0.46] for Monkey 1, [0.36, 0.42] for Monkey 2; Low vs Low in Fig. 4B). Similarly, the dynamics of unit-level high task information populations were distinct from those for unit-level low task information populations (95% CI: [0.38, 0.47] for Monkey 1, [0.33, 0.42] for Monkey 2; Low vs High in Fig. 4B). The dynamics of unit-level high task information populations were also significantly more correlated than that of randomly selected populations (95% CI: [0.62, 0.72] for Monkey 1, [0.55, 0.67] for Monkey 2; Rand in Fig. 4B). Units with high task information had correlated dynamics across higher CC modes, providing evidence that the correlated dynamics were unlikely to be driven by trivial correlations captured by the first few CC modes (Fig. 4C). The link between task information and dynamics is also consistent with the observation that spatial distributions of population decoding accuracy (see Fig. 2G, H) and the units with similar dynamics were qualitatively (Fig. S9A) and quantitatively (Fig. S9B) similar.

If the link between task information and correlated dynamics is indeed at the unit-level, we would expect that neural populations selected based on task information would have more similar dynamics than neural populations selected based on other factors, including recording site. To test this, we randomly selected neural populations composed of units recorded at the same recording site and compared their dynamics. We found that the dynamics similarity of neural populations selected based on recording site was significantly lower than neural populations selected based on task information (95% CI: [0.65, 0.75] for Monkey 1, [0.55, 0.72] for Monkey 2; Within site in Fig. 4B, C). We also controlled for the possibility that pseudo-population analyses could obscure shared dynamics (e.g., by failing to capture correlations related to non-task relevant signals that vary across sessions). We replicated our unit-level group analyses restricted to simultaneously recorded populations (which were also from the same recording site) and found that high task information populations consistently had significantly higher dynamics similarity than other sub-populations (no CI overlap across populations at any recording site; see *Statistical significance estimation*; Fig. 4D).

The correlations we observed within high task-information neural populations were significantly larger during the task than during inter-trial intervals, a commonly used a lower-bound estimate on population correlations (see *CCA lower bound estimation*; Fig. 4B). Low task-information populations, in contrast, remained near the lower bounds, consistent with a link between task-related activity and correlated neural dynamics. Correlated dynamics within the unit-level high task-information population, however, were unlikely to be a trivial result of the relationship between task information and neural dynamics. As noted above, empirically, we found that CCA correlation and decoding accuracy were weakly correlated when neural populations were composed of randomly selected units. Yet, the dynamics similarities observed within our high task information sub-populations exceed those expected from this correlation alone (Fig. S8D, E). In fact, randomly selected neural populations that were decoding accuracy-matched to the high task information populations all had lower CCA correlations than the high task information sub-populations (95% CI: [0.66, 0.76] for Monkey 1, [0.61, 0.72] for Monkey 2; Acc match in Fig. 4B; S8F, G). We also found similar results when analyzing neural populations composed of only task-responsive units (see *Task-responsive unit estimation*), suggesting that task *information*, not only responsiveness, was the primary driver of dynamics similarity (Fig. S7H-I). Together, our results indicated that correlated dynamics between two neural populations were primarily driven by highly task-informative units.

### Spatially distributed populations of units have similar task representations

Our previous analysis revealed that neural populations were more likely to have similar dynamics when they contain units with high task information. Yet, it remains unclear if those similar dynamics also reflect a shared task representation. If two neural populations have similar task representations, a decoder trained on one will generalize to the other. We therefore analyzed patterns of decoding generalization across neural populations defined by task information. Specifically, we aligned the dynamics of each neural population with CCA, then trained a decoder on one neural population and tested it on the other (Fig. 5A) (Gallego et al., 2020; Safaie et al., 2023).

We found that decoders trained on the aligned dynamics of unit-level high task information populations consistently generalized to other unit-level high task information populations (95% CI: [98.7, 100.5] for Monkey 1, [96.7, 100.2] for Monkey 2; High vs High in Fig. 5B), even though the units themselves were spatially distributed across frontal motor cortices (see Fig. 4D). This was not the case between two unit-level low task information populations, which performed at or near chance level (95% CI: [10.8, 21.5] for Monkey 1, [10.1, 18.8] for Monkey 2; Low vs Low in Fig. 5B). We also found that decoders trained on the aligned dynamics from low task information populations generalized above chance to high task information populations, though performance was variable across populations (95% CI: [54.9, 83.6] for Monkey 1, [49.3, 77.6] for Monkey 2; Low vs High in Fig. 5B). This is consistent with a distribution of task information (see Fig. 2) where even the neural populations with the lowest relative task information had enough information to train decoders that performed above chance level. These shared task representations did not appear to be a trivial result of correlated dynamics, which can be independent conceptually (Fig. S10A-C). Similar to our within-population decoding analyses, we found a weak correlation between CCA correlation and aligned decoding accuracy when neural populations were composed of randomly selected units (R2: 0.22 for Monkey 1, 0.27 for Monkey 2; Fig S10D). Yet, the dynamics similarities observed within our high task information sub-populations exceed that expected from this correlation alone (Fig. S10D, E).

Our results suggest that neural populations in frontal motor cortices contain a mixture of distinct and shared dynamics, and the exact proportion of distinct and shared dynamics vary across populations. High task information populations primarily exhibited shared dynamics driven by a similar task representation. Other neural populations had a higher proportion of distinct dynamics potentially from participating in other motor related computations and processes.

## Discussion

Limits in our ability to sample neural population activity have made it difficult to fully resolve the structure of task information in frontal motor cortices. We used precise targeting of high-density laminar microelectrode arrays to better map spatiotemporal dynamics of individual neurons and neural populations across frontal motor cortices areas. Decoding analyses revealed variability across both the cortical surface and depths, with significantly more task information at some locations than others and no consistent pattern in depth relationships across cortical locations. Population dynamics analysis also revealed variability in the temporal structure of task-related population activity across the cortical surface. Neurons that were most informative about the task were distributed in space, yet also had activity patterns that evolved coherently in time. Our findings highlight the complex spatiotemporal structure of frontal motor cortex and reveal potential links between heterogeneity in task information and population dynamics. Our largescale neuron-resolution maps provide new perspectives on the seeming contradictions across studies mapping motor cortices at different scales, while also highlighting the need to consider spatiotemporal complexity for translational applications like BCIs.

### Spatiotemporal heterogeneity in task decoding and dynamics

Our results extend previous findings that single-unit tuning notably varies across neurons (Schieber and Hibbard, 1993; Georgopoulos et al., 2007) to neural populations distributed across frontal motor cortices. We found that task information was spatially distributed (Fig. 2G, H) but primarily clustered in units recorded at the same cortical surface location (Fig. S9). Task information was similarly distributed across all cortical depths, where the depth with the most task information varied across cortical surface locations (Fig. 2I, J). Our depth analysis was limited by the complicated sulcal topography of frontal motor cortex. We inserted probes perpendicular to the implanted grid surface, preventing us from confidently estimating the location of layer boundaries. Despite these limitations, our findings are consistent with previous work suggesting minimal spatial organization in motor cortical activity at the level of single neurons.

Population-level perspectives that consider how neurons coordinate to control movements, meanwhile, have suggested a more unified picture of motor cortices. Our results add to this literature by revealing significant variability in population dynamics across frontal cortices. Caudal areas near the central sulcus are known to be predominantly involved in movement execution compared to rostral areas (Riehle and Requin, 1989; Cisek and Kalaska, 2005; Ouchi et al., 2025). Consistent with this, we found a weak correlation between rostral-caudal location and population dynamics (Fig. 3F, G). However, patterns in population dynamics were more strongly linked to the spatial distribution of task information across the cortical surface (Fig. 3I, J), potentially facilitated by strong coordination specifically among the highly task modulated units (Fig. 4). These results motivate future investigations into co-occurring neural dynamics related to non-task relevant activity, which will be best resolved by widespread simultaneous recordings. Furthermore, our results open questions about how spatially distributed neurons may coordinate to produce movements and highlights the need to integrate neuron-level and population-level approaches to fully characterize the neural representations of movement.

Our analyses suggest that subsets of neurons, identifiable from their unit-level properties (e.g., high task information), tightly coordinate during the task consistent with recent mechanisms of population coordination (Perich et al., 2018; Semedo et al., 2019; Veuthey et al., 2020). Yet, our results also highlight that this sub-population captures neural dynamics shared across all neurons in motor cortices, as revealed by patterns of decoding generalization (Fig. 5). The presence of shared task-relevant dynamics across all neurons is consistent with the ability to decode movement information through population recordings with non-targeted sampling (e.g., fixed geometry multi-electrode arrays) (Georgopoulos et al., 1986; Churchland et al., 2012). Neurons sharing some task-relevant dynamics are also likely related to observations that stable latent cortical dynamics underlie consistent behavior (Gallego et al., 2020). Past studies focused on the collective dynamics of all neurons (Churchland et al., 2012; Elsayed et al., 2016; Gallego et al., 2018, 2020), but the contributions of individual neurons to dynamics will likely need to be further considered. Our results suggest that the strength of the shared dynamics between two neural populations will vary, predicting that neurons that participate in other computations would coordinate differently. This generates testable hypotheses for future studies given the many forms of information about movement variables observed in motor cortices (e.g. planning vs. execution (Riehle and Requin, 1989; Cisek and Kalaska, 2005); arm vs. eye movement (Cisek and Kalaska, 2002; Pesaran et al., 2010; Ouchi et al., 2025); or movement variables encoded in different coordinate systems (Evarts, 1968; Fetz, 1994; Kakei et al., 1999; Sergio et al., 2005). While focusing on target information, our results add new insights into the variable temporal structure observed in single-neuron activity (Churchland and Shenoy, 2007) by suggesting that the similarity of dynamics between distinct neural populations is driven by different combinations of a shared task-relevant component.

### Spatially-distributed functional networks

Spatially focal recordings suggest that performing well-learned movements recruits consistent subsets of neurons with precise timing (Rubino et al., 2006; Peters et al., 2014; Best et al., 2016). Patterns in spike-timing between neurons also show that the spatiotemporal structure of these recruited neural populations are task-specific, suggesting they form a functionally defined network (Moore et al., 2024). Consistent with this, we found a subset of spatially distributed units encoded significant amounts of task information and did so with highly correlated dynamics, which were distinct from the dynamics of other neural populations (Fig. 4B, C). Our results extend previous findings by suggesting that neurons participating in these functionally defined networks are spatially distributed across cortical areas and depths during well-learned center-out reaching tasks in macaques. Whether and how the functional network structure varies across species and behavioral tasks is an important question for future studies.

The potential relationship between population dynamics and connections between neurons (Sussillo et al., 2015) raises questions about the structural basis of functional networks. Anatomical studies found long-range connections across frontal motor cortex supporting cortico-cortical communication (Gatter and Powell, 1978; Huntley and Jones, 1991; Ninomiya et al., 2019). Functional studies suggest that subcortical structures contribute to functional network coordination (Ghanayim et al., 2025; Ramot et al., 2025). Both communication pathways would provide mechanisms for spatially distributed neurons to form functional networks, and future research investigating links to anatomical connectivity will further improve our understanding of motor representations.

If well-learned motor tasks recruit specialized functional networks, this raises questions about whether and how neurons flexibly contribute to multiple computations and, if so, how multiple networks form. Different movements, or muscle activation patterns, seem to display distinct correlation patterns between neurons (Moore et al., 2024), consistent with the possibility that distinct computations involve distinct networks. Our analyses focused on neural activity during movement execution, but frontal motor cortices also exhibit distinct, but linked, population-level activity during movement planning (Elsayed et al., 2016). Our results make the testable prediction that these computations might be implemented by distinct, spatially distributed functional networks. Recent work also finds that motor learning shapes the spatiotemporal recruitment of individual neurons (Peters et al., 2014). Such plasticity mechanisms may contribute to forming functionally-defined networks, though significant work remains to investigate how multiple functional networks coexist and coordinate during complex motor tasks.

### Implications for BCI

Our results also highlight the need to consider the spatial distribution of movement information when targeting implants for BCIs. Current BCIs have primarily targeted mesoscale microelectrode array implants (with fixed geometry) based on functional cortical mapping (Hochberg et al., 2006; Collinger et al., 2013; Willett et al., 2021; Kunigk et al., 2024). Our results (Fig. 2) show that the functional organization of task information may be at a smaller scale than accessible by functional mapping techniques like functional MRI and intracortical micro-stimulation. These mapping approaches, then, may be limited in their utility for targeting spatially-precise, flexible neural implants (Jun et al., 2017; Musk and Neuralink, 2019; Yang et al., 2019). Optimizing task information decoding for BCIs may require new mesoscale and causal mapping techniques to precisely target implants to functionally-relevant networks, or dense non-targeted recording strategies paired with computational methods to identify functionally-relevant networks within large datasets (e.g., feature selection).

Improved mapping approaches may also be needed to target coordinated neural populations. Our results revealed that population temporal dynamics differ across distinct neural populations. A mismatch between the temporal structure of neural activity with those expected by a BCI can reduce performance (Sadtler et al., 2014; Oby et al., 2019, 2025), suggesting that BCIs may benefit from targeting recordings to neural populations that share common dynamics. Successfully mapping temporal dynamics will also likely help ensure robust BCI performance over long timescales in the presence of recording instabilities. Recent approaches to maintain performance when neural measurements change specifically leverage the temporal structure of neural population activity (Degenhart et al., 2020; Karpowicz et al., 2025). Our results suggest that functionally defined networks have distinct dynamics, which could obscure the ideal temporal dynamics for maintaining stable BCI task performance. Methods to identify functional networks and shared task-relevant dynamics among neural populations will likely improve longitudinal BCI decoding approaches.

Together, our study reveals the spatiotemporal complexity of neural activity patterns across frontal motor cortical areas at the level of individual neurons and populations. New insights into the spatiotemporal distribution of motor representations in the brain will be critical for leveraging advanced neural probe technologies for BCIs.

## Acknowledgments

We would like to thank the Washington National Primate Research Behavioral Management team for their consultations on animal training. This work was supported in part by a National Center for Advancing Translational Sciences of the National Institutes of Health fellowship (TL1 TR002318, R.A.C.), a Nakajima Foundation fellowship (T.O.), a postdoctoral fellow award from Weill Neurohub (L.R.S.), a Simons Collaboration for the Global Brain Pilot award (898220, A.L.O.), the National Science Foundation Harnessing the Data Revolution Program (2117997, A.L.O., H.F.), the National Institute of Neurological Disorders and Stroke (NIH R01 NS134634 and NIH UF1NS126485, A.L.O.), and the NIH Office of Research Infrastructure Programs (P51 OD010425).

## Supplementary Materials

**Figure S1.**
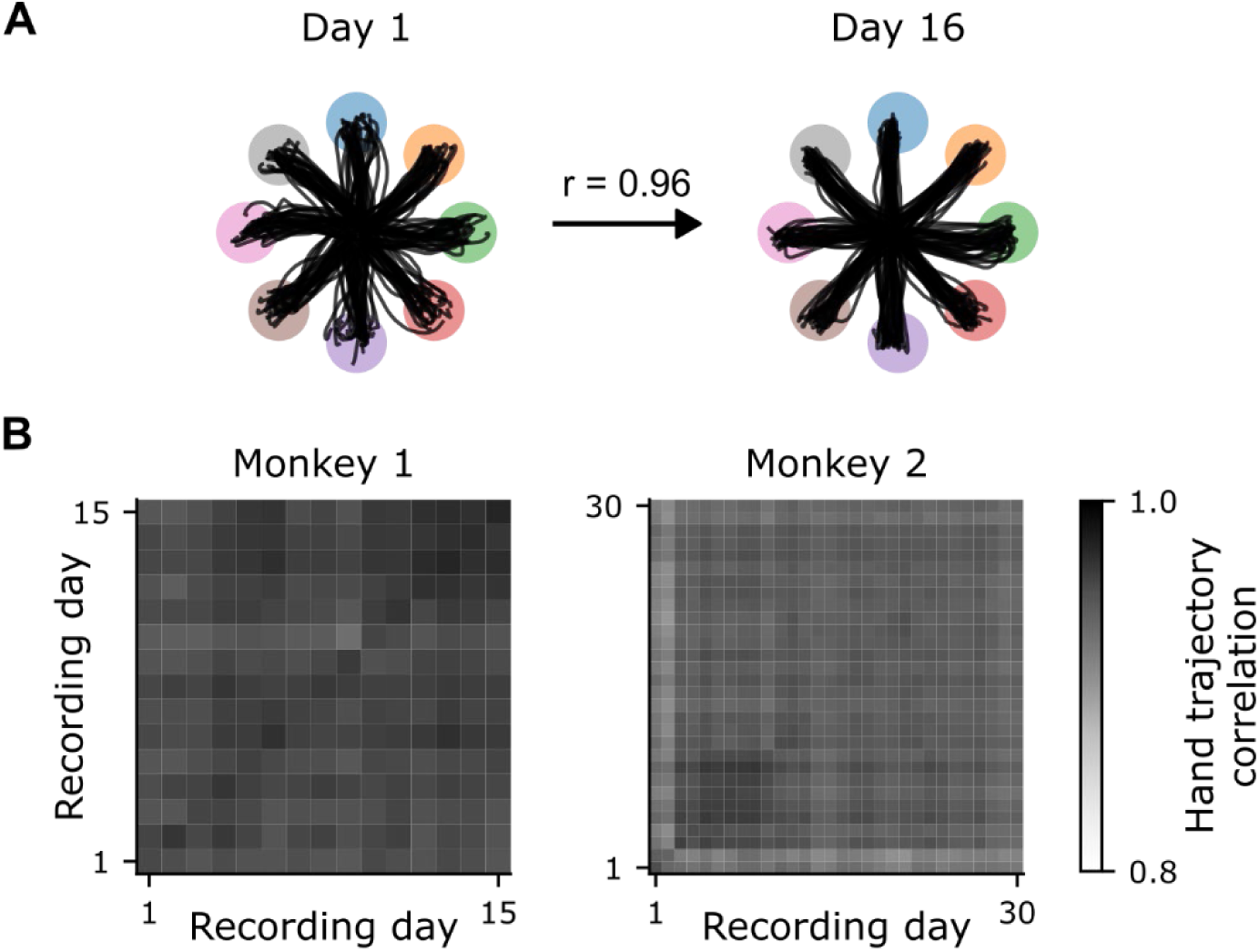
Three-dimensional hand trajectories are correlated across days. **(A)** Individual trial cursor trajectories for Monkey 1 on two different example days (day 1, left; day 16, right), and the average cursor trajectory correlation between those days (r = 0.96, n = 240). **(B)** Average hand trajectory correlation across all combinations of day pairs for each monkey. Only 15 days are reported for Monkey 1 because of a technical error where 3D kinematics were not recorded for a single day.

**Figure S2.**
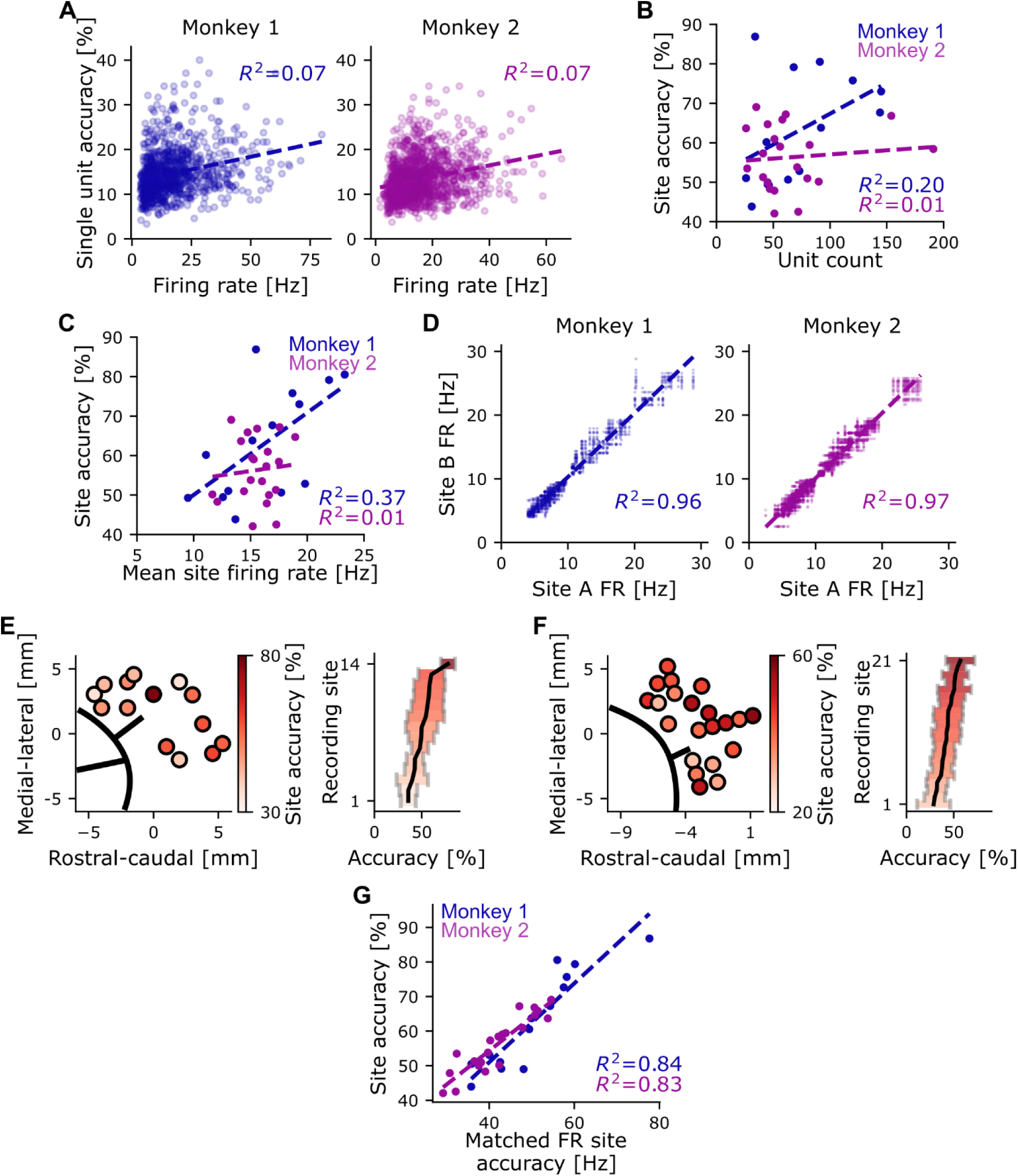
Decoding accuracy variability is not explained by firing rate. **(A)** Relationship between single-unit decoding accuracy and the average firing rate of each recorded unit (N=1,022 for monkey 1; N=1,441 for monkey 2). **(B)** Relationship between number of recorded units and site accuracy for each recording site (N=14 for monkey 1; N=21 for monkey 2). **(C)** Relationship between the average firing rate of all recorded units at each site and the site accuracy for each recording site (N=14 for monkey 1; N=21 for monkey 2). **(D)** Relationship of unit firing rates for all pairs of firing rate-matched recording site populations (referred to as A, B) (N=1456 for Monkey 1; N=3360 for Monkey 2). **(E)** Left: Distribution of firing rate-matched population decoding accuracy across the cortical surface (site accuracy) for monkey 1. The black lines represent the arcuate sulcus. Right: Population decoding accuracy across recording sites, sorted by the mean decoding accuracy. Black lines indicate the mean; the shaded region represents the 95% CI. Colors indicate the site decoding accuracy (color bar) for both right and left panels. **(F)** Same as (E) but for Monkey 2. **(G)** Relationship of decoding accuracies estimated in the firing rate-matched condition and the non-matched condition (N=14 for monkey 1; N=21 for monkey 2). For all panels, dashed lines represent the linear regression.

**Figure S3.**
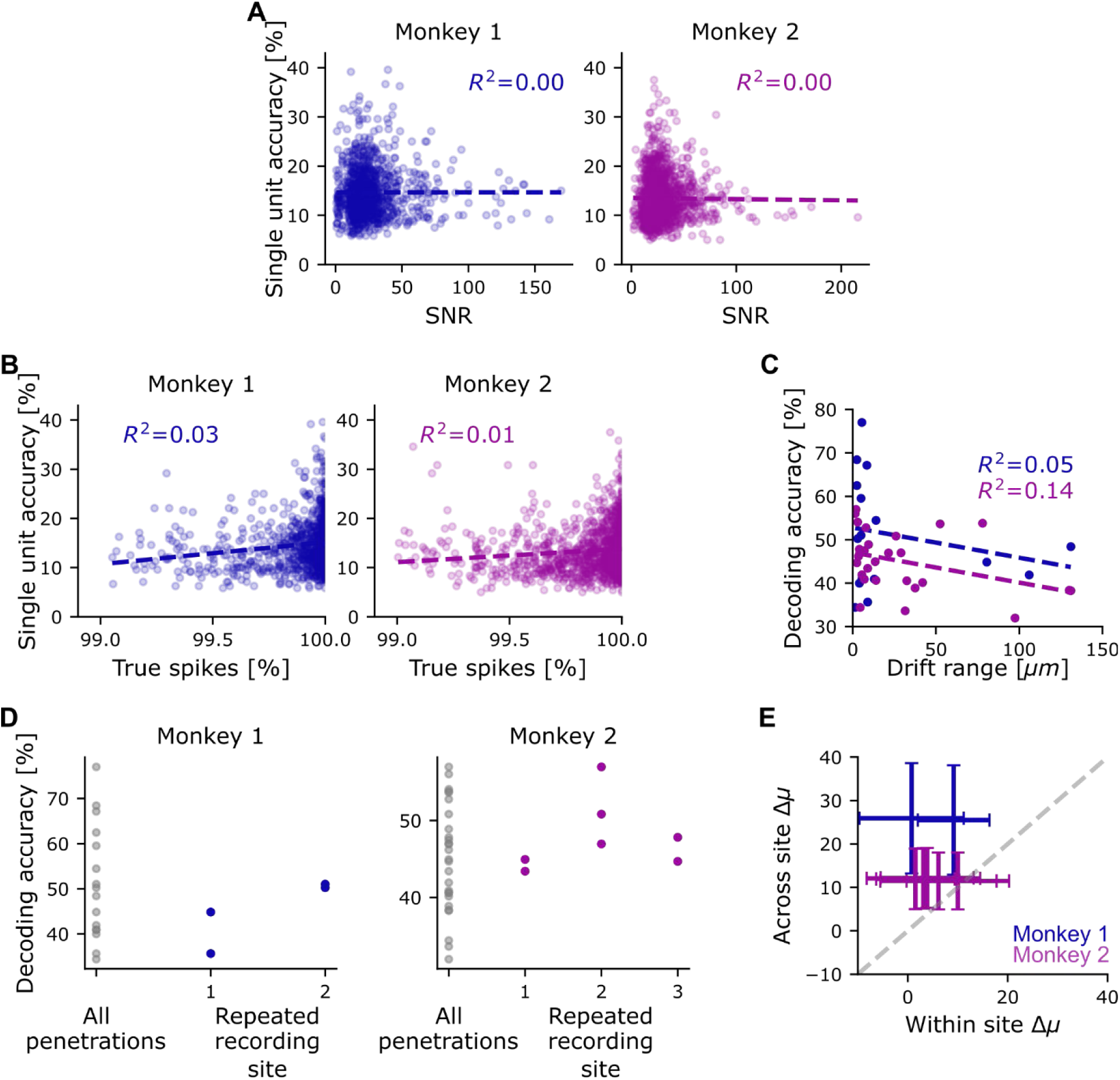
Decoding accuracy variability is not explained by recording quality. **(A)** Relationship between single-unit decoding accuracy and the signal-to-noise ratio for each unit (see *Unit quality metrics*) **(B)** Relationship between the single-unit decoding accuracy and true spike percentage based on refractory period violations (see *Unit quality metrics*). **(C)** Relationship between the drift range computed by Kilosort 4 and decoding accuracy for each probe insertion. **(D)** Average decoding accuracy for each penetration at a recording site. Decoding accuracy from each repeated probe penetration (blue/purple) next to the decoding accuracy from all other probe penetrations for each monkey (gray). **(E)** The mean decoding difference for repeated penetrations compared to the mean decoding difference across sites. Error bars represent the standard deviation across unit group resampling (see *Statistical analysis across multiple penetrations*). The gray dashed line represents the identity line. Note: All colored dashed lines represent the linear regression fit.

**Figure S4.**
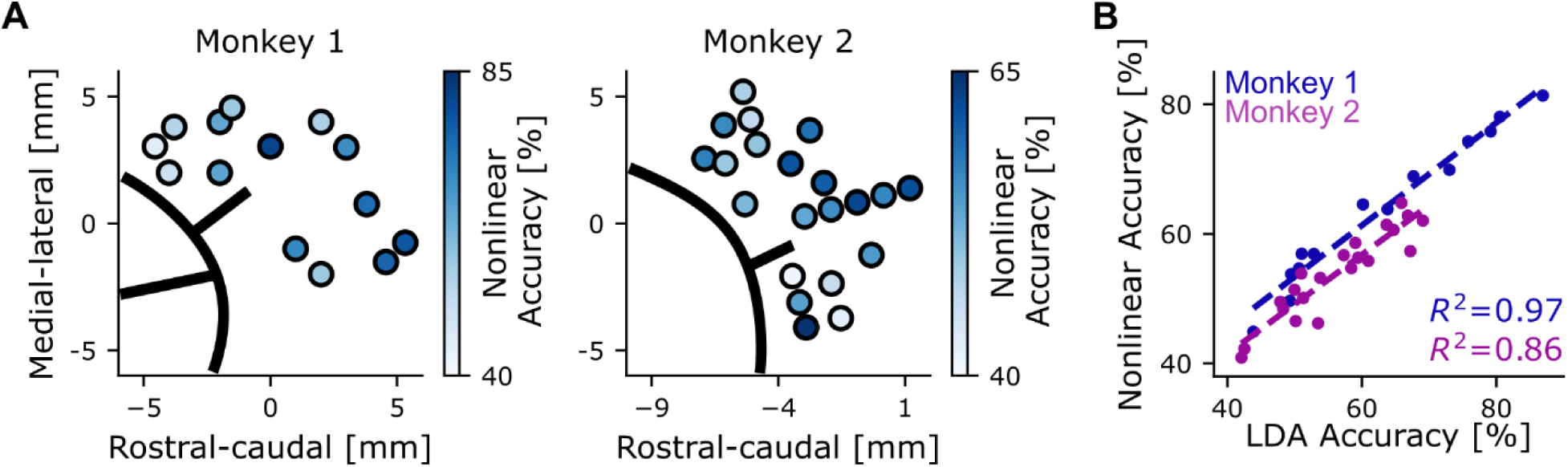
Decoding accuracy is correlated when computed with either linear or nonlinear methods. **(A)** Site accuracy computed with a nonlinear decoding algorithm (see *Nonlinear decoding analysis*). **(B)** Relationship between nonlinear decoding accuracy and linear decoding accuracy (see *Linear decoding analysis*). Dashed lines represent the linear regression fit.

**Figure S5.**
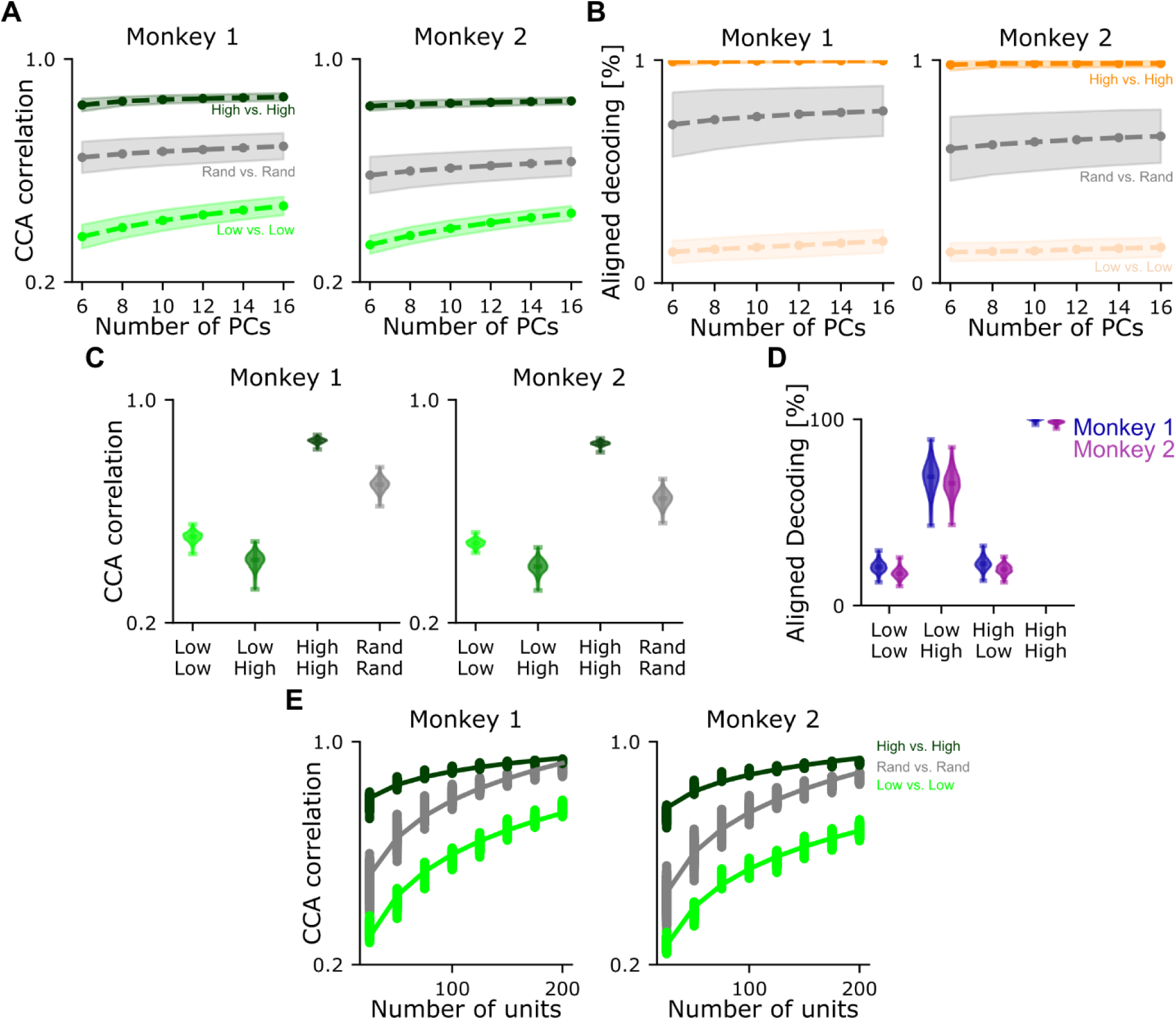
Dynamic correlations are specific to high task information populations across a range of principal components (PC). **(A)** CCA correlation for task neural populations across a range of principal components (PC). Dashed lines represent the mean across random unit sampling, and the shaded regions represent the 95% CI. **(B)** Same as (A) but for aligned decoding. **(C)** CCA correlation for task responsive neural populations while controlling for the explained variance. Each distribution represents the variability from all random unit resampling centered on the mean. **(D)** Formatted the same as (C), but for aligned decoding. **(E)** CCA correlation for task responsive neural populations across a range of neural population sizes. Solid lines indicate an exponential fit to the data. Each dot represents the CCA correlation from random unit resampling using the number of units indicated by the x-axis. For all panels, high and low decoding accuracy units were defined as the 100 units with the highest and lowest single-unit decoding accuracy, respectively (see *Shared neural population dynamics between task-tuned populations* and Fig. 4A).

**Figure S6.**
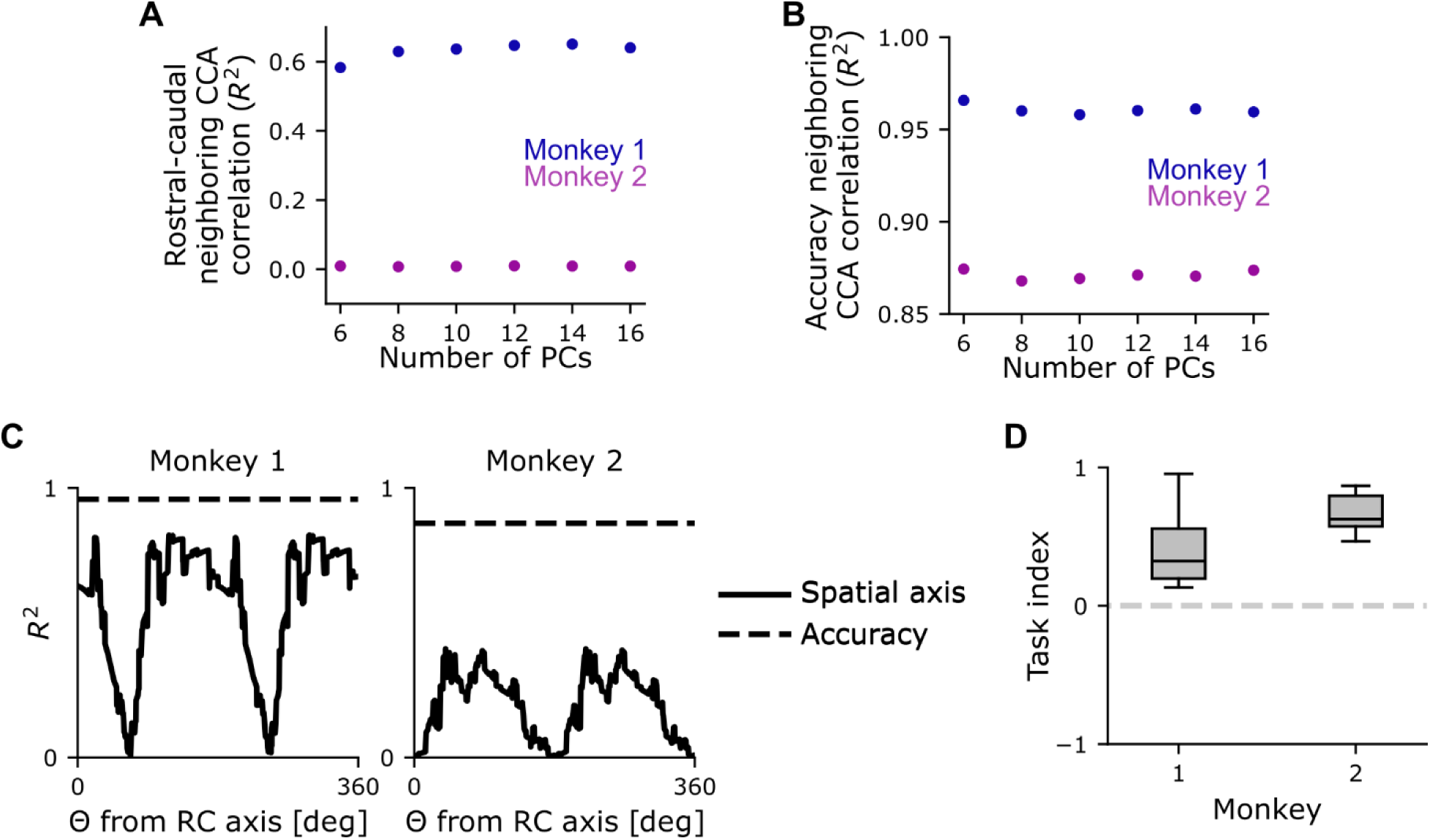
Spatial organization results are consistent across a reasonable range of principal components (PC) and across different spatial axes. **(A)** *R*^2^ of the relationship between neighboring CCA correlation and rostral-caudal position of recording sites across a range of principal components (PC). **(B)** Same as (A) except with recording sites organized by decoding accuracy. **(C)** Relationship of the *R*^2^ from (A) except when computed with recording sites organized along a variety of different axis relative to the rostral-caudal (RC) axis (solid line) using 10 PCs. Dashed line represents the *R*^2^ from (B) with 10 PCs. **(D)** Task index computed for each axis (C). Positive values indicate that decoding accuracy predicts similar dynamics better than spatial position (see *Neural population dynamics between recording sites*). Dashed line represents a task index of 0 where decoding accuracy and spatial axis predict correlated dynamics similarly well.

**Figure S7:**
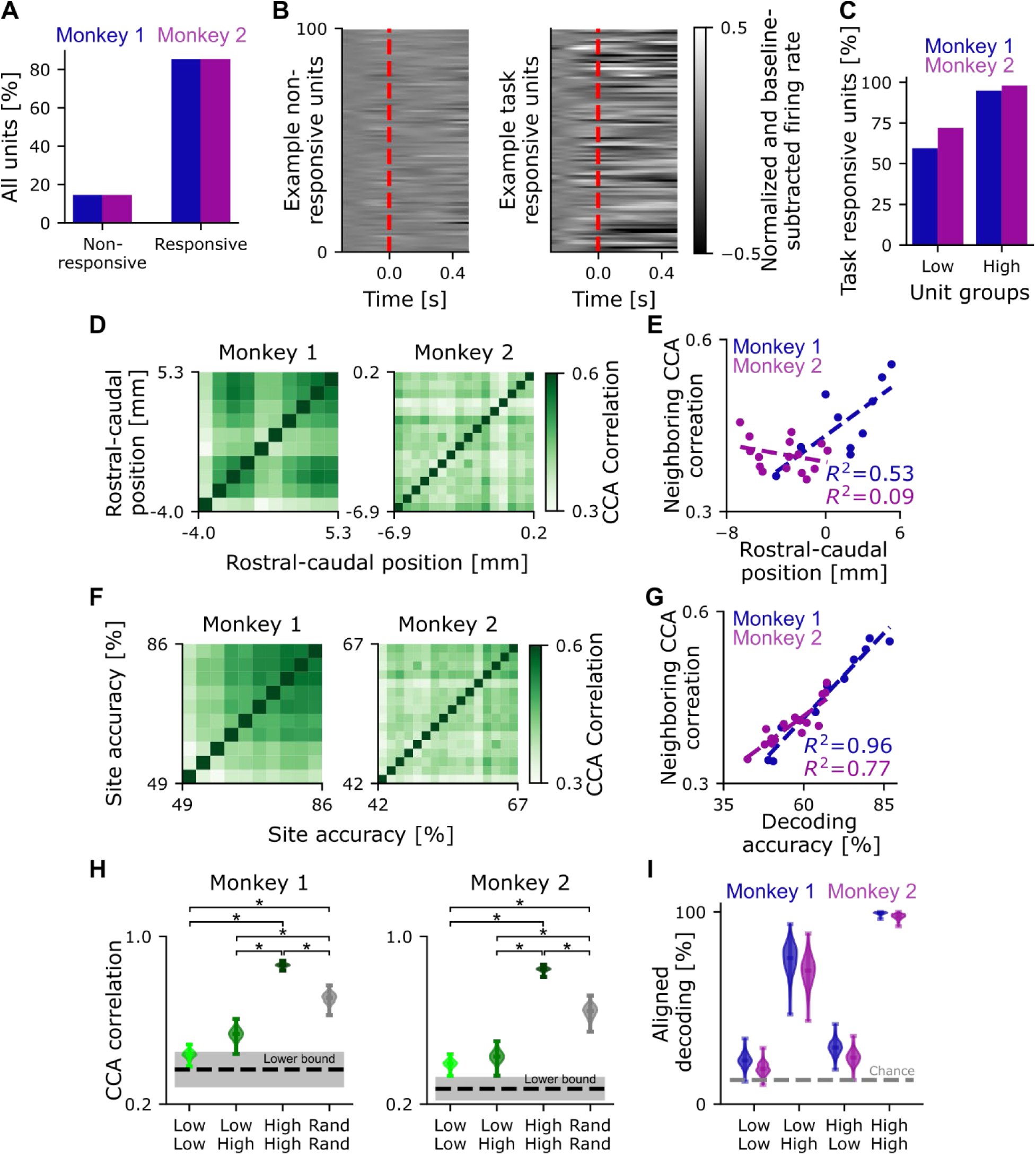
Dynamic correlations are specific to high task information populations when analysis is restricted to task-responsive units. **(A)** Percentage of all recorded units identified as not task responsive and task responsive for monkey 1 (blue) and monkey 2 (purple). **(B)** Trial averaged and baseline-subtracted firing rate as a function of time of 100 randomly selected example units from Monkey 1 that were identified as not task-responsive (left) or task-responsive (right; see *Task-responsive unit estimation*). Firing rates were time-aligned to movement onset, indicated by the red dashed line. **(C)** Percent of task-responsive units in the high and low task information unit groups from Fig. 4 that included non-responsive units. **(D)** Heatmap of CCA correlation between neural populations of only task-responsive units organized by rostral-caudal position for Monkey 1 (left) and Monkey 2 (right). Each pixel represents a pair of neural populations (N=45 for monkey 1; N=120 for monkey 2). **(E)** Neighboring CCA correlation based on rostral-caudal position for each distinct neural population (see *Neural population dynamics between recording sites*) for monkey 1 (blue) and monkey 2 (pink). Each dot represents a recording site (N=10 for monkey 1; N=16 for monkey 2). The dashed line represents the linear regression. **(F)** Heatmap of CCA correlation between distinct neural populations organized by the amount of task information for Monkey 1 (left) and Monkey 2 (right). Format as in (B). **(G)** Neighboring CCA correlation based on task information for each distinct neural population, formatted as in (C). **(H)** CCA correlation between distinct neural populations of task-responsive units defined by task information (Low and High) or randomly selected (Rand) for monkey 1 (left) and monkey 2 (right). The distribution for each comparison represents the variability from different random unit selections (N=1000). Each distribution depicts the CCA correlation results from all random task-responsive unit selections and is centered on the mean. The black dashed line represents the lower bound estimated from the inter-trial interval activity of task-responsive units (see *CCA lower bound estimation*) and the gray shading represents the 95% CI across random unit selections. **(I)** Aligned decoding for each neural population comparison. The distribution represents the variability of aligned decoding from different random task-responsive unit selection (N=1000). Each distribution depicts the aligned decoding results from all random task-responsive unit selections and is centered on the mean. The gray dashed line represents chance-level decoding.

**Figure S8.**
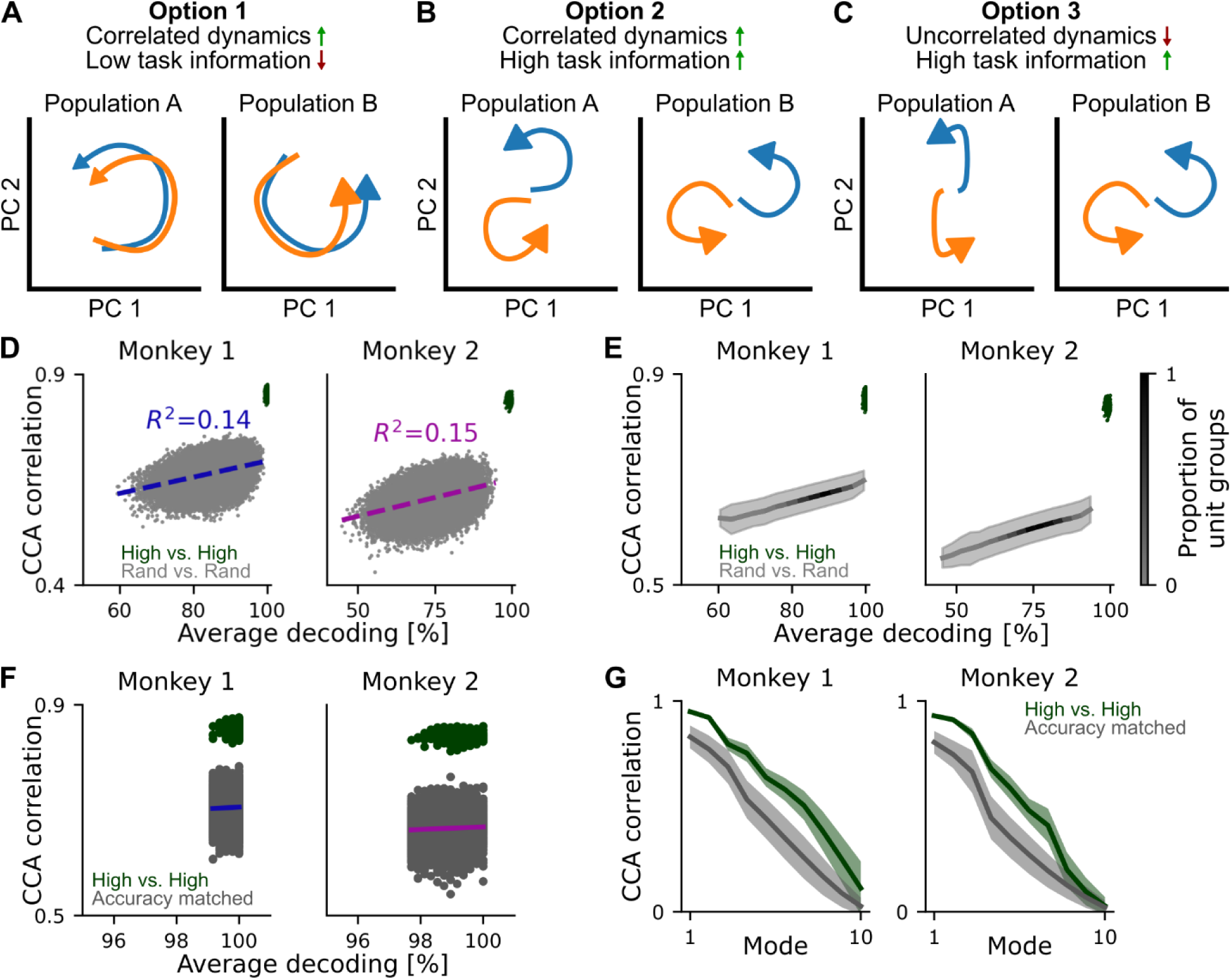
Correlated dynamics are not necessarily solely a consequence of high task information. **(A)** Conceptual schematic diagram depicting correlated dynamics with low task information. Left: Population dynamics in PC space for neural population A. Each color represents the trajectory for a different task condition (i.e. target direction). Right: Same as Left, but for neural population B, composed of different units. **(B)** Same as (A) but for correlated dynamics with high task information **(C)** Same as (A) but for uncorrelated dynamics with high task information **(D)** CCA correlation between two distinct neural populations as a function of their average decoding accuracy. Gray dots represent a comparison between neural populations composed of randomly selected units (N=100,000). Green dots represent a comparison between neural populations composed of high task information units (N=1,000). Each dot is from different random unit sampling. The dashed line represents the linear regression. **(E)** Same as D except representing the distribution of results from unit groups. The solid line represents the mean, and the shaded region represents ±1σ. The gradient represents where units were distributed along the x-axis, normalized to the maximum number of groups. **(F)** Same as D but with random unit selection restricted to produce populations that match the range of decoding accuracies observed in the high-task information subpopulations (N=6536 for Monkey 1 and N=5562 for Monkey 2). The colored line represents the linear regression of the randomly selected neural populations. **(G)** CCA correlation as a function of CC mode. The shaded regions represent the 95% confidence interval for each mode across random unit selections.

**Figure S9.**
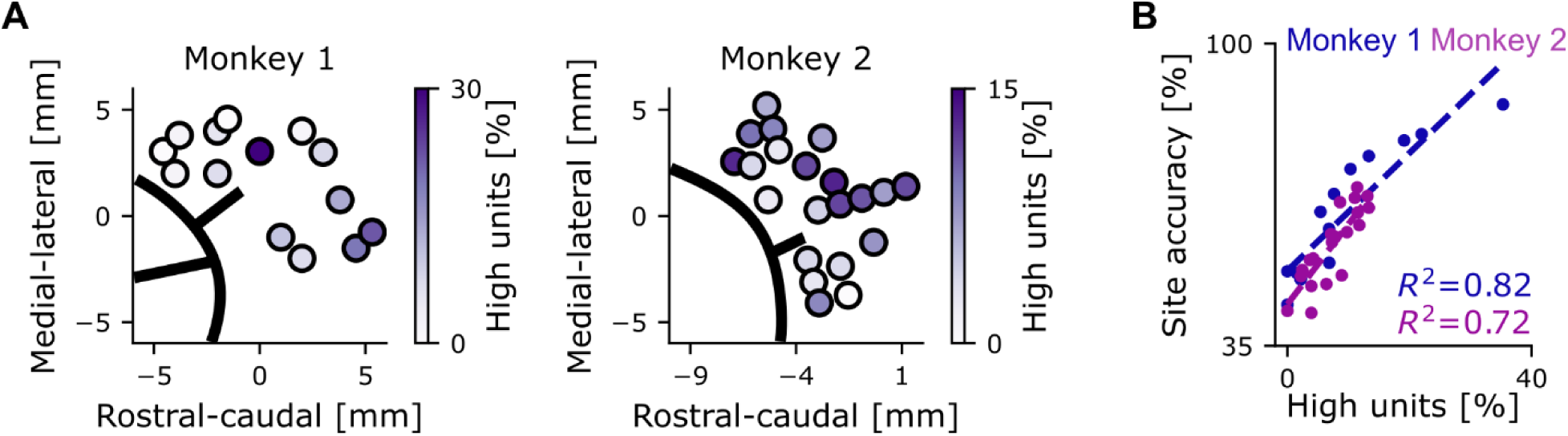
The spatial distribution of the high task information unit locations is correlated with the spatial distribution of population decoding accuracy. **(A)** Spatial distribution of the recording location for high task information units for Monkey 1 (left) and Monkey 2 (right). Each circle represents a recording site, and the x-y position indicates the coordinates within the chamber. The color at each recording site represents the percentage of units with high task information relative to the total number of units measured at each recording site. **(B)** Population decoding accuracy for a recording site (same analysis as Fig. 2G, H) as a function of the percentage of units with high task information recorded at that site for monkey 1 (blue) and monkey 2 (pink). Each data point represents a recording site (N=14 for monkey 1; N=21 for monkey 2); dashed lines represent a linear regression (R^2^ = 0.82, p < 0.0001 for monkey 1; R^2^ = 0.72, p < 0.0001 for monkey 1).

**Figure S10:**
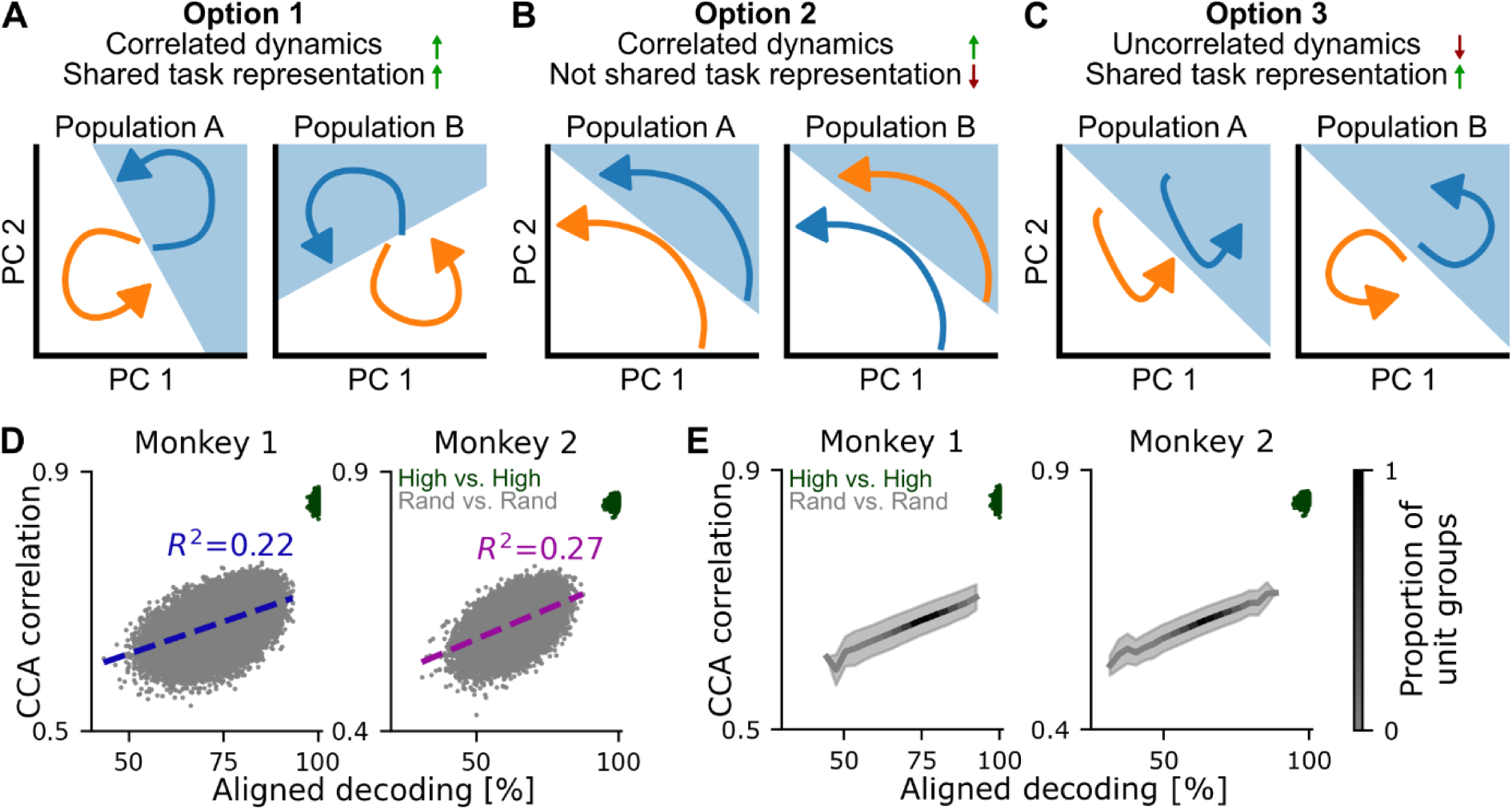
A shared task representation is not required by correlated dynamics. **(A)** Diagram depicting correlated dynamics with a shared task representation. Left: Population dynamics in PC space for neural population A. Each color represents the trajectory for a different task condition (i.e. target direction). Right: Same as Left, but for neural population B, composed of different units. The blue shaded background represents a decision boundary between task conditions trained on Population A, then tested on Population B after alignment between Population A and B. **(B)** Same as A but for correlated dynamics without a shared task representation. **(C)** Same as A but for uncorrelated dynamics with a shared task representation. **(D)** CCA correlation between two neural populations as a function of their aligned decoding. Gray dots represent neural populations composed of randomly selected units (N=100,000). Green dots represent neural populations composed of high task information units (N=1,000). Each dot is from different random unit sampling. The dashed line represents the linear regression. **(E)** Same as D except representing the distribution of results from unit groups. The solid line represents the mean, and the shaded region represents ±1σ. The gradient represents where units were distributed along the x-axis, normalized to the maximum number of groups.

## References

Amirikian B, Georgopoulos AP (2003) Modular organization of directionally tuned cells in the motor cortex: Is there a short-range order? Proc Natl Acad Sci 100:12474–12479.

Asanuma H, Rosén I (1972) Topographical organization of cortical efferent zones projecting to distal forelimb muscles in the monkey. Exp Brain Res 14:243–256.

Banerjee A, Dean HL, Pesaran B (2010) A Likelihood Method for Computing Selection Times in Spiking and Local Field Potential Activity. J Neurophysiol 104:3705–3720.

Best MD, Suminski AJ, Takahashi K, Brown KA, Hatsopoulos NG (2016) Spatio-Temporal Patterning in Primary Motor Cortex at Movement Onset. Cereb Cortex 27 Available at: https://academic.oup.com/cercor/article/doi/10.1093/cercor/bhv327/3056241.

Chehade NG, Gharbawie OA (2023) Motor actions are spatially organized in motor and dorsal premotor cortex. eLife 12:e83196.

Churchland MM, Cunningham JP, Kaufman MT, Foster JD, Nuyujukian P, Ryu SI, Shenoy KV (2012) Neural population dynamics during reaching. Nature 487:51–56.

Churchland MM, Shenoy KV (2007) Temporal Complexity and Heterogeneity of Single-Neuron Activity in Premotor and Motor Cortex. J Neurophysiol 97:4235–4257.

Cisek P, Kalaska JF (2002) Modest Gaze-Related Discharge Modulation in Monkey Dorsal Premotor Cortex During a Reaching Task Performed With Free Fixation. J Neurophysiol 88:1064–1072.

Cisek P, Kalaska JF (2005) Neural Correlates of Reaching Decisions in Dorsal Premotor Cortex: Specification of Multiple Direction Choices and Final Selection of Action. Neuron 45:801– 814.

Collinger JL, Wodlinger B, Downey JE, Wang W, Tyler-Kabara EC, Weber DJ, McMorland AJ, Velliste M, Boninger ML, Schwartz AB (2013) High-performance neuroprosthetic control by an individual with tetraplegia. The Lancet 381:557–564.

Dadarlat MC, Canfield RA, Orsborn AL (2023) Neural Plasticity in Sensorimotor Brain–Machine Interfaces. Annu Rev Biomed Eng 25:51–76.

Degenhart AD, Bishop WE, Oby ER, Tyler-Kabara EC, Chase SM, Batista AP, Yu BM (2020) Stabilization of a brain–computer interface via the alignment of low-dimensional spaces of neural activity. Nat Biomed Eng 4:672–685.

Dum R, Strick P (2002) Motor areas in the frontal lobe of the primate. Physiol Behav 77:677–682.

Elsayed GF, Lara AH, Kaufman MT, Churchland MM, Cunningham JP (2016) Reorganization between preparatory and movement population responses in motor cortex. Nat Commun 7:13239.

Evarts EV (1968) Relation of pyramidal tract activity to force exerted during voluntary movement. J Neurophysiol 31:14–27.

Fetz EE (1994) Are movement parameters recognizably coded in the activity of single neurons? In: Movement Control, 1st ed. (Cordo P, Harnad S, eds), pp 77–88. Cambridge University Press. Available at: https://www.cambridge.org/core/product/identifier/CBO9780511529788A013/type/book_part.

Gallego JA, Perich MG, Chowdhury RH, Solla SA, Miller LE (2020) Long-term stability of cortical population dynamics underlying consistent behavior. Nat Neurosci 23:260–270.

Gallego JA, Perich MG, Naufel SN, Ethier C, Solla SA, Miller LE (2018) Cortical population activity within a preserved neural manifold underlies multiple motor behaviors. Nat Commun 9:4233.

Gatter KC, Powell TPS (1978) The intrinsic connections of the cortex of area 4 of the monkey. Brain 101:513–541.

Georgopoulos A, Kalaska J, Caminiti R, Massey J (1982) On the relations between the direction of two-dimensional arm movements and cell discharge in primate motor cortex. J Neurosci 2:1527–1537.

Georgopoulos AP, Merchant H, Naselaris T, Amirikian B (2007) Mapping of the preferred direction in the motor cortex. Proc Natl Acad Sci 104:11068–11072.

Georgopoulos AP, Schwartz AB, Kettner RE (1986) Neuronal Population Coding of Movement Direction. Science 233:1416–1419.

Ghanayim A, Benisty H, Cohen Rimon A, Schwartz S, Dabdoob S, Lifshitz S, Talmon R, Schiller J (2025) VTA projections to M1 are essential for reorganization of layer 2-3 network dynamics underlying motor learning. Nat Commun 16 Available at: https://www.nature.com/articles/s41467-024-55317-4.

Hochberg LR, Serruya MD, Friehs GM, Mukand JA, Saleh M, Caplan AH, Branner A, Chen D, Penn RD, Donoghue JP (2006) Neuronal ensemble control of prosthetic devices by a human with tetraplegia. Nature 442:164–171.

Huntley GW, Jones EG (1991) Relationship of intrinsic connections to forelimb movement representations in monkey motor cortex: a correlative anatomic and physiological study. J Neurophysiol 66:390–413.

International Brain Laboratory (2024) Spike sorting pipeline for the International Brain Laboratory. :18689452 Bytes Available at: https://figshare.com/articles/online_resource/Spike_sorting_pipeline_for_the_International_Brain_Laboratory/19705522/4.

Jun JJ et al. (2017) Fully integrated silicon probes for high-density recording of neural activity. Nature 551:232–236.

Kakei S, Hoffman DS, Strick PL (1999) Muscle and Movement Representations in the Primary Motor Cortex. Science 285:2136–2139.

Karpowicz BM, Ali YH, Wimalasena LN, Sedler AR, Keshtkaran MR, Bodkin K, Ma X, Rubin DB, Williams ZM, Cash SS, Hochberg LR, Miller LE, Pandarinath C (2025) Stabilizing brain-computer interfaces through alignment of latent dynamics. Nat Commun 16:4662.

Kaufman MT, Churchland MM, Ryu SI, Shenoy KV (2014) Cortical activity in the null space: permitting preparation without movement. Nat Neurosci 17:440–448.

Kunigk NG, Schone HR, Gontier C, Hockeimer W, Tortolani AF, Hatsopoulos NG, Downey JE, Chase SM, Boninger ML, Dekleva BD, Collinger JL (2024) Motor somatotopy impacts imagery strategy success in human intracortical brain-computer interfaces. Available at: http://medrxiv.org/lookup/doi/10.1101/2024.08.01.24311180.

Lara AH, Cunningham JP, Churchland MM (2018) Different population dynamics in the supplementary motor area and motor cortex during reaching. Nat Commun 9:2754.

Mollazadeh M, Aggarwal V, Davidson AG, Law AJ, Thakor NV, Schieber MH (2011) Spatiotemporal Variation of Multiple Neurophysiological Signals in the Primary Motor Cortex during Dexterous Reach-to-Grasp Movements. J Neurosci 31:15531–15543.

Moore DD, MacLean JN, Walker JD, Hatsopoulos NG (2024) A dynamic subset of network interactions underlies tuning to natural movements in marmoset sensorimotor cortex. Nat Commun 15:10517.

Musk E, Neuralink (2019) An Integrated Brain-Machine Interface Platform With Thousands of Channels. J Med Internet Res 21:e16194.

Ninomiya T, Inoue K, Hoshi E, Takada M (2019) Layer specificity of inputs from supplementary motor area and dorsal premotor cortex to primary motor cortex in macaque monkeys. Sci Rep 9:18230.

Oby ER, Degenhart AD, Grigsby EM, Motiwala A, McClain NT, Marino PJ, Yu BM, Batista AP (2025) Dynamical constraints on neural population activity. Nat Neurosci.

Oby ER, Golub MD, Hennig JA, Degenhart AD, Tyler-Kabara EC, Yu BM, Chase SM, Batista AP (2019) New neural activity patterns emerge with long-term learning. Proc Natl Acad Sci 116:15210–15215.

Ouchi T, Scholl LR, Rajeswaran P, Canfield RA, Smith LI, Orsborn AL (2025) Mapping eye, arm, and reward information in frontal motor cortices using electrocorticography in non-human primates. J Neurosci:e1536242025.

Pachitariu M, Sridhar S, Pennington J, Stringer C (2024) Spike sorting with Kilosort4. Nat Methods 21:914–921.

Pandarinath C, O’Shea DJ, Collins J, Jozefowicz R, Stavisky SD, Kao JC, Trautmann EM, Kaufman MT, Ryu SI, Hochberg LR, Henderson JM, Shenoy KV, Abbott LF, Sussillo D (2018) Inferring single-trial neural population dynamics using sequential auto-encoders. Nat Methods 15:805–815.

Park MC, Belhaj-Saïf A, Gordon M, Cheney PD (2001) Consistent Features in the Forelimb Representation of Primary Motor Cortex in Rhesus Macaques. J Neurosci 21:2784–2792.

Pedregosa F, Varoquaux G, Gramfort A, Michel V, Thirion B, Grisel O, Blondel M, Prettenhofer P, Weiss R, Dubourg V, Vanderplas J, Passos A, Cournapeau D, Brucher M, Perrot M, Duchesnay É (2011) Scikit-learn: Machine Learning in Python. J Mach Learn Res 12:2825– 2830.

Perich MG, Gallego JA, Miller LE (2018) A Neural Population Mechanism for Rapid Learning. Neuron 100:964–976.e7.

Pesaran B, Nelson MJ, Andersen RA (2010) A Relative Position Code for Saccades in Dorsal Premotor Cortex. J Neurosci 30:6527–6537.

Peters AJ, Chen SX, Komiyama T (2014) Emergence of reproducible spatiotemporal activity during motor learning. Nature 510:263–267.

Ramot A, Taschbach FH, Yang YC, Hu Y, Chen Q, Morales BC, Wang XC, Wu A, Tye KM, Benna MK, Komiyama T (2025) Motor learning refines thalamic influence on motor cortex. Nature 643:725–734.

Riehle A, Requin J (1989) Monkey primary motor and premotor cortex: single-cell activity related to prior information about direction and extent of an intended movement. J Neurophysiol 61:534–549.

Rubino D, Robbins KA, Hatsopoulos NG (2006) Propagating waves mediate information transfer in the motor cortex. Nat Neurosci 9:1549–1557.

Sadtler PT, Quick KM, Golub MD, Chase SM, Ryu SI, Tyler-Kabara EC, Yu BM, Batista AP (2014) Neural constraints on learning. Nature 512:423–426.

Safaie M, Chang JC, Park J, Miller LE, Dudman JT, Perich MG, Gallego JA (2023) Preserved neural dynamics across animals performing similar behaviour. Nature 623:765–771.

Schieber MH, Hibbard LS (1993) How Somatotopic Is the Motor Cortex Hand Area? Science 261:489–492.

Semedo JD, Zandvakili A, Machens CK, Yu BM, Kohn A (2019) Cortical Areas Interact through a Communication Subspace. Neuron 102:249–259.e4.

Sergio LE, Hamel-Pâquet C, Kalaska JF (2005) Motor Cortex Neural Correlates of Output Kinematics and Kinetics During Isometric-Force and Arm-Reaching Tasks. J Neurophysiol 94:2353–2378.

Silversmith DB, Abiri R, Hardy NF, Natraj N, Tu-Chan A, Chang EF, Ganguly K (2020) Plug-and-play control of a brain-computer interface through neural map stabilization. Nat Biotechnol 39:326–335.

Sussillo D, Churchland MM, Kaufman MT, Shenoy KV (2015) A neural network that finds a naturalistic solution for the production of muscle activity. Nat Neurosci 18:1025–1033.

Veuthey TL, Derosier K, Kondapavulur S, Ganguly K (2020) Single-trial cross-area neural population dynamics during long-term skill learning. Nat Commun 11:4057.

Willett FR, Avansino DT, Hochberg LR, Henderson JM, Shenoy KV (2021) High-performance brain-to-text communication via handwriting. Nature 593:249–254.

Williams JJ, Rouse AG, Thongpang S, Williams JC, Moran DW (2013) Differentiating closed-loop cortical intention from rest: building an asynchronous electrocorticographic BCI. J Neural Eng 10:046001.

Yang X, Zhou T, Zwang TJ, Hong G, Zhao Y, Viveros RD, Fu T-M, Gao T, Lieber CM (2019) Bioinspired neuron-like electronics. Nat Mater 18:510–517.

